# Protein-Driven Copper Redox Regulation: Uncovering the Role of Disulphide Bonds and Allosteric Modulation

**DOI:** 10.1101/2025.08.20.671251

**Authors:** Rebecca Sternke-Hoffmann, Chang Liu, Xue Wang, Hegne Pupart, Xun Sun, Jan Gui-Hyon Dreiser, Peep Palumaa, Qinghua Liao, Matthias Krack, Jinghui Luo

## Abstract

Copper plays essential roles in enzymatic activity, redox reactions, and cellular signalling, but becomes toxic when redox homeostasis is disrupted. While Cu(II) reduction is commonly attributed to unfolded or amyloid proteins, here we show that the well-folded plasma protein human serum albumin (HSA) intrinsically reduces Cu(II) to Cu(I) in the absence of external reductants. Using X-ray absorption spectroscopy (XAS), small-angle X-ray scattering (SAXS), circular dichroism (CD) and computational modelling (QM/MM and DFT), we identify a redox mechanism involving the disulphide bond Cys392-Cys438 in domain III of HSA. Cu binding at the high-affinity ATCUN motif triggers conformational changes that expose this disulphide bond, enabling thiol-mediated electron transfer and Cu(I) formation. Chelation with tetrathiomolybdate (TTM) impairs this reduction by restricting access to the reactive disulphide site. Comparative analysis with other globular proteins reveals that Cu reduction requires both accessible disulphide motifs and a native folded structure. Simulations and spectroscopy of SOD1 confirm that disulphide cleavage enhances Cu-thiolate interaction, supporting a generalizable two-site redox mechanism. These findings reveal a previously unrecognized mode of protein-mediated copper reduction and suggest broader physiological roles for disulphide-regulated redox switching in metal homeostasis.

## Introduction

Living organisms depend on various transition metals for proper functioning, and copper is a prime example. Copper plays a crucial role in enzymatic catalysis, intracellular signalling, and redox balance^1^.

Copper’s redox versatility enables it to participate in single-electron transfers, primarily existing in two oxidation states: Cu(I) and Cu(II)^2^. Although this property is vital for enzyme functions, it also contributes to oxidative stress through the formation of hydroxyl radicals through the Haber-Weiss reaction, potentially damaging biomolecules^3^. To mitigate such risks, organisms have developed a sophisticated regulatory system that collectively maintains copper homeostasis through specific proteins^4,5^.

Intracellularly, Cu(I) predominates in the reducing environment maintained by glutathione (GSH) and cysteine^6^. In contrast, Cu(II) is stabilised in extracellular fluids, especially within high-affinity protein-binding sites^7^. The redox conversion between these two states is central to copper’s biological activity and toxicity. Cysteine residues, in particular, play key roles in both binding and redox conversion, due to their thiol chemistry^8–10^. Copper coordination preferences vary by oxidation state. Cu(I), a d^10^ cation, exhibits minimal stabilization energy of the ligand field and prefers coordination with soft ligands such as His, Cys and Met^11,12^. Cu(II), a d^9^ cation, adopts square planar, square pyramidal, or axially distorted octahedral geometries due to Jahn-Teller distortion^13^. Ligand preferences align with the theory of hard and soft acids and bases (HSAB), as demonstrated by cyclen-based ligands that removed Cu(II) from amyloid-*β* (A*β*), while sulphur-containing analogues preferentially bind Cu(I)^14,15^.

Previous studies have primarily linked Cu(II) reduction to amyloid and intrinsically disordered proteins, implicating histidine, tryptophan, and cysteine residues in redox activity^16–25^. Amyloid-β (Aβ), for instance, can reduce Cu(II) to Cu(I)^24^, which may exacerbate oxidative stress and contribute to neurotoxicity. Some studies propose that amyloid proteins facilitate the transport of copper across cellular membranes, but pathological aggregation may exacerbate Cu(II) reduction and oxidative stress^26^. However, the physiological relevance of this process remains debated, as Aβ concentrations *in vivo* are far lower than those for plasma proteins such as human serum albumin (HSA)^27,28^. In the bloodstream, copper is primarily bound to metalloproteins, such as ceruloplasmin, while a small labile fraction interacts with proteins, mainly HSA, amino acids, and peptides^29–31^.

Disruptions in copper regulation contribute to disease pathogenesis. Wilson’s disease (WD) and Menke’s disease (MD) arise from mutations in the copper transporters ATP7B and ATP7A, respectively, leading to copper mislocalization^32^. In WD, excessive copper accumulation damages the liver and brain, while in MD, copper deficiency disrupts essential enzymatic functions. Furthermore, Amyotrophic Lateral Sclerosis (ALS) is linked to mutations in the metalloprotein Cu,Zn-superoxide dismutase (SOD1), which impair copper binding, contribute to protein aggregation and thereby hinder the neutralisation of free radicals^32^. Understanding copper homeostasis is also critical for therapeutic development, such as optimizing metal chelators like tetrathiomolybdate (TTM).

Here, we investigate whether well-folded globular proteins also possess intrinsic Cu(II)-reducing activity in the absence of external reductants. We focus on HSA, the most abundant plasma protein and a major copper carrier in the bloodstream^33^. HSA contains a high-affinity Cu(II)-binding site known as the ATCUN (Amino-Terminal Cu- and Ni-binding) motif, as well as additional metal binding regions and 17 disulphide bonds^34^. HSA and its ATCUN motif DAH exhibit a strong Cu(II) affinity with K_*d*_ of 9.55 x 10^−14^ and 2.00 × 10^−14^, respectively^35, 36^. Beyond HSA, ATCUN motifs are found in diverse proteins across biological compartments^37,38^. While previous work suggested that Cu(I) formation on HSA requires external reductants like ascorbate, we now demonstrate that HSA alone can reduce Cu(II) to Cu(I) under physiological conditions.

To elucidate this unexpected redox behaviour, we combine X-ray absorption spectroscopy (XAS), small-angle X-ray scattering (SAXS), circular dichroism (CD), and computational simulations (MD, QM/MM, and DFT). we compare the redox efficiency between HSA and three different ATCUN motif peptides: DAH (from HSA), DTHFPI (from hepcidin)^39,40^, and MEHFPGP (from semax)^41,42^. While previous studies suggest that these peptides display strong Cu(II) affinity^35,41,43–46^, direct competition experiments indicate comparable binding strengths, with DAH exhibiting the lowest dissociation constant^36^. We identify a previously unrecognised redox-active disulphide bond (Cys392-Cys438) in HSA’s domain III distant from the ATCUN motif. Cu binding at the ATCUN motif induces allosteric rearrangements that expose this site, facilitating copper reduction via thiol coordination.

To assess the generality of this mechanism, we extend our analysis to other globular proteins (including β-lactoglobulin, insulin, lysozyme, and the metalloprotein SOD1). While only HSA shows strong Cu(II)-reducing activity in its native fold, SOD1 also reveals comparable disulphide-gated Cu redox switch upon bond cleavage. These findings suggest that intrinsic redox regulation via conformational gating of disulphide bonds may be a broader property of globular proteins than previously appreciated.

## Results

### HSA Reduces Copper via a Mechanism Independent of Its High-affinity ATCUN motif

HSA, a key copper transporter, can bind up to five copper ions^48^. Three primary binding sites have been identified in its N-terminal domain. These include the ATCUN motif (Asp-Ala-His), a multi-metal site at the IA-IIA interface, and a reduced cysteine at position 34, which are indicated on the crystal structure (Fig. 1a).

**Figure 1.**
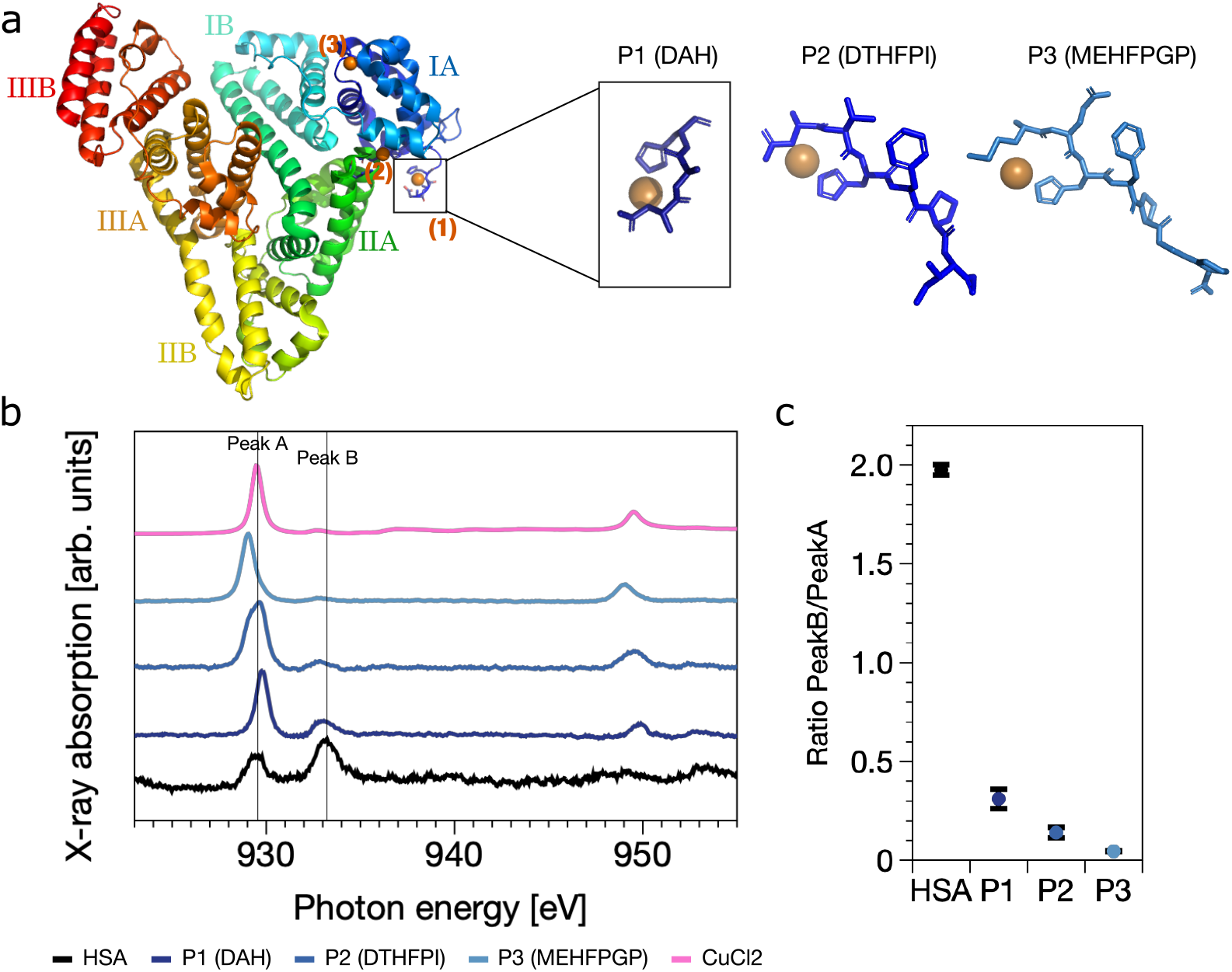
HSA reduces Cu(II) to Cu(I) via mechanism distinct from ATCUN motif. (a) Structural representations of human serum albumin (HSA, PDB ID: 7WLF^47^) and the three ATCUN peptides: DAH (P1), DTHFPI (P2), and MEHFPGP (P3), generated via AlphaFold. In HSA, the primary copper-binding sites are highlighted in orange and located around domain I. (b) Soft X-ray absorption spectra recorded at the Cu L_2,3_-edges for HSA (black), ATCUN motif peptides P1 (dark blue), P2 (blue), and P3 (light blue), incubated with CuCl_2_ (1:2 molar ratio) at pH 7.4. CuCl_2_ alone is shown in pink. Peak A (Cu(II), 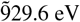) and B (Cu(I), 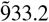 eV) are indicated with vertical lines. (c) Ratio of integrated Peak B to Peak A areas, quantifying the extent of Cu(I) formation. HSA shows a substantially higher Cu(I)/Cu(II) ratio than any ATCUN peptide, suggesting that the ATCUN motif does not drive the observed reduction. Hard X-ray absorption data at the Cu K-edge (Fig. S1 confirm the presence of Cu(I) in HSA and reveal a distinct coordination geometry compared to ascorbate-reduced Cu(I), supporting the involvement of a different copper-binding site.

To investigate the role of HSA’s high-affinity ATCUN motif in copper reduction, we compared full-length HSA with three peptides containing canonical ATCUN motifs - DAH (P1), DTHFPI (P2), and MEHFPGP (P3) - using soft X-ray absorption spectroscopy (XAS). Upon incubation with CuCl_2_ (1:2 molar ratio (protein:CuCl_2_), the Cu L-edge spectra of HSA displayed two key features: Peak A (at 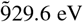), characteristic of Cu(II), and Peak B (at 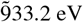), indicative of monovalent Cu(I) (Fig. 1b). In contrast, the ATCUN peptides showed only Peak A with minimal or no Peak B signal (Fig. 1b and c, Fig. S1). This comparison indicated that Cu(II) is not reduced by the ATCUN motif alone and that the reduction observed in HSA occurs through a distinct mechanism.

Hard X-ray spectroscopy at the Cu K-edge confirmed the presence of Cu(I) in HSA and revealed that its coordination geometry differs significantly from the two-coordinate linear Cu(I) species formed by ascorbate reduction (Fig. S2). This suggests that Cu(I) is reduced at a site outside the canonical high-affinity binding motifs and adopts a likely four-coordinate geometry^49,50^.

### Chelation Reveal Allosteric Modulation of Copper Reduction in HSA

To explore whether Cu(II) reduction in HSA involves allosteric mechanisms or specific structural regions, we examined the impact of tetrathiomolybdate (TTM), a high-affinity copper-chelator, on copper redox behaviour and protein conformation. TTM is commonly used in the treatment of Wilson’s disease and cancer therapy, as it effectively depletes bioavailable intracellular copper^52,53^. XAS spectra of Cu-TTM complexes in solution and with ATCUN peptides resembled Cu(I)(Fig. 2a), indicating that TTM stabilises the reduced state, consistent with its strong Cu(I) binding capability. When mixed with ATCUN peptides, TTM led to a nearly complete loss of the Cu(II) signal, suggesting that peptide-bound Cu is easily accessible for chelation and reduction. The shift from cupric copper in the absence of chelating agent toward cuprous copper in the presence of TTM was particularly evident for peptide P3 (see Fig. S1).

**Figure 2.**
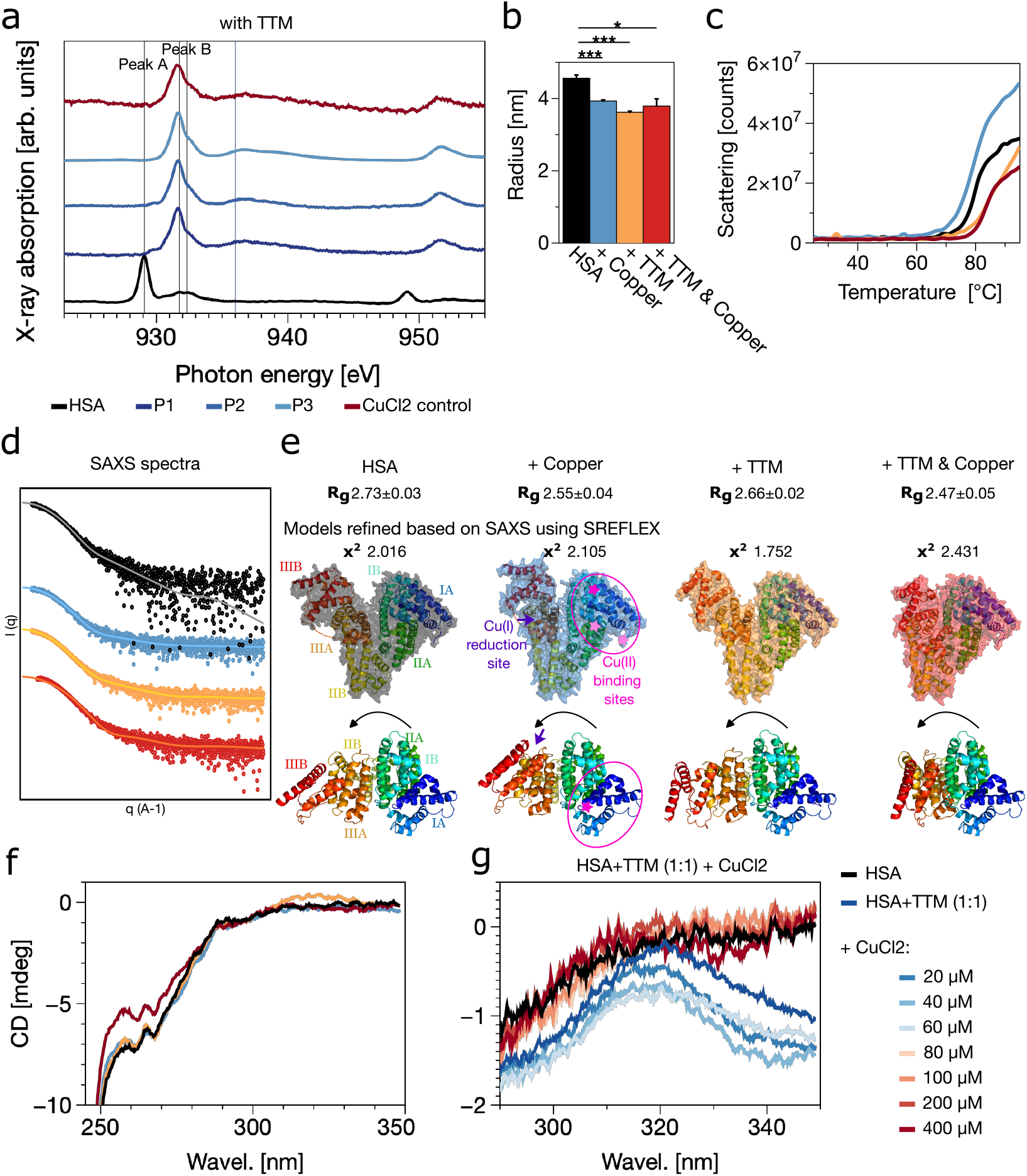
Copper chelation reveal allosteric regulation of Cu(II) reduction in HSA. (a) Soft X-ray absorption spectra at the Cu L_2,3_-edges for 100 µM HSA (black) and three ATCUN peptides (DAH (P1 dark blue), DTHFPI (P2, blue), and MEHFPGP (P3, light blue)) incubated with CuCl_2_ (200 µM) and TTM (100 µM) at pH 7.4. While Cu(II) signals are suppressed in peptides (spectra resemble Cu-TTM control), HSA retains a prominent Cu(II) peak, suggesting TTM interferes with reduction in HSA. (b) Hydrodynamic radius of HSA measured by DLS. Copper and TTM both lead to compaction. (c) DSF-derived thermal unfolding profiles show increased thermal stability upon TTM binding, consistent with structural stabilisation. (d) SAXS profiles (black: HSA; blue: +CuCl_2_; orange: +TTM; red: +CuCl_2_+TTM) with model fits (lighter colours) and (e) SAXS-derived models (via SREFELX, 4LA0^51^) show compaction localised to domain IIIA/IIIB (red/orange), while canonical Cu-binding sites in domain I (marked with pink) remain structurally unchanged. (f) Near-UV CD spectra reveal changes in aromatic residue environments, especially in Cu-TTM-HSA complex. (g) CD signal around 320 nm (indicative of disulphide formation) appears upon TTM binding but diminishes upon Cu titration, suggesting competition between TTM and Cu for redox-active cysteines.

However, in HSA, TTM had the opposite effect: Cu(II) was retained, and Cu(I) formation was suppressed. This indicates that TTM interferes with the reduction process, either by limiting Cu mobility, shielding redox-active sites, or altering HSA’s structure. To support this, dynamic light scattering (DLS) and differential scanning fluorimetry (DSF), circular dichroism (CD) measurements confirmed that TTM binding induces structural changes in HSA, compacting and stabilizing the protein (Fig. 2b, c). SAXS analysis (see Fig. S3) with SREFELX modelling localised these conformational changes in domain IIIA and IIIB, which are distal to the Cu-binding ATCUN motif in domain I (Fig. 2d, e). However, increasing the CuCl_2_ and TTM concentrations (≥1:3) led to destabilisation, aggregation and partially unstructured conformations (as shown in Figures S4, S5, S6, S7). These structural changes were absent in domain I, where the canonical Cu-binding sites are located, suggesting an allosteric transmission of Cu binding events to more distal redox-active regions.

Circular dichroism further confirmed local structural rearrangements and perturbations of the disulphide bond upon TTM and copper binding (Fig. 2f, g). A slight perturbation in the local environment of phenylalanine and tyrosine residues could be observed within the Cu-TTM-HSA complex. TTM alone induced a signal consistent with disulphide formation (Fig. S8), which was abolished by titration of Cu(II), suggesting a dynamic interplay between disulphide status and copper redox activity. In general, allosteric changes induced by TTM might obstruct access to redox-active disulphide site and thereby inhibiting copper reduction.

These findings support the idea that copper binding initiates allosteric structural rearrangements that propagate to remote regions containing disulphide bonds.

### Disulphide Bond Cys392-Cys438 is a Likely Redox-Active Site for Cu(II) Reduction in HSA

We next analysed disulphide bond geometry using SAXS-refined models to identify potential redox-active sites. Among the 17 disulphide bonds in HSA, most of which are buried in a stable α-helical regions, Cys392 and Cys438 stood out in SAXS-refined models (Fig. 3). The Cα-Cα distance between these residues increased from 4.1 Å in the apo-state to 7.2 Å with copper and 8.4 Å with both copper and TTM. These values exceed typical disulphide bond length^54^, suggesting destabilisation or partial cleavage.

**Figure 3.**
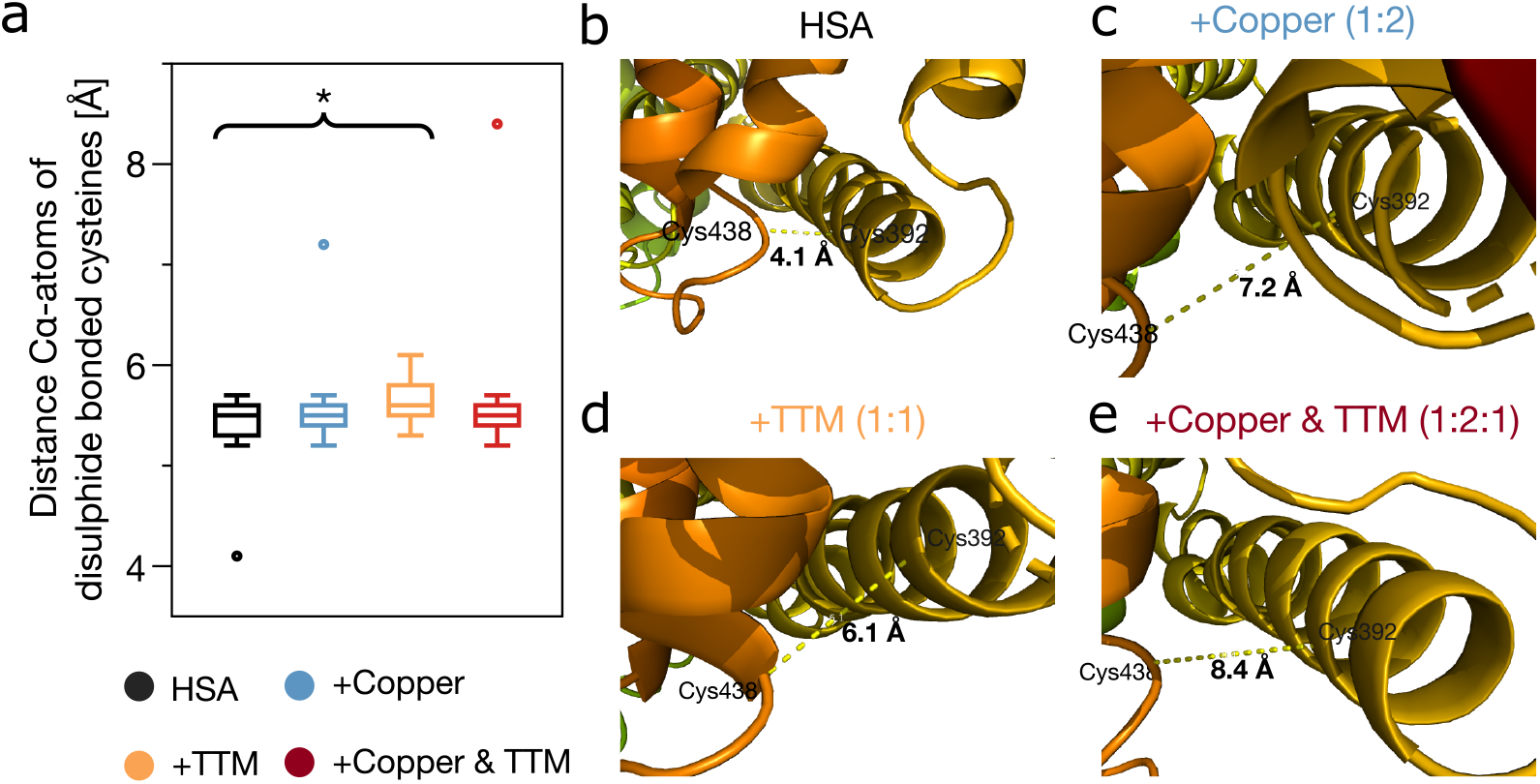
SAXS modelling reveals domain III disulphide Cys392-Cys438 is structurally perturbed during copper redox modulation. (a) Quantification of average Cα-Cα distances across all 17 disulphide bonds in HSA based on SAXS-refined models under different conditions: apo HSA (black), with CuCl_2_ (blue), with TTM (orange), and with both CuCl_2_ and TTM (red). While most disulphide bond distances remain stable, a significant expansion is observed in the TTM-treated sample (p=0.0026). The primary outlier was the Cys392–Cys438 bond, indicating selective disulphide perturbation. (b-e) Structural visualisation of the Cys392-Cys438 disulphide bond from SAXS-refined HSA models. The Cα-Cα distance increases progressively from the native state (b, 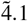 Å) to Copper (c, 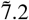 Å), TTM (d, 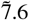 Å) and Copper+TTM (e, 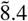 Å), exceeding typical disulphide geometries and supporting redox-active function.

This bond is located in domain III, which undergoes the most prominent conformational rearrangements in SAXS. No comparable distance changes were observed in other disulphide binds. This site is also structurally positioned near His440 and Arg445, potentially facilitating interaction with copper. This structural evidence, combined with CD and SAXS data showing compaction and disulphide rearrangement in domain III strongly suggests that Cys393-Cys438 is involved in Cu(II) reduction. The domain’s flexibility and proximity to known fatty acids and metal-binding sites make it a plausible reduction site, likely regulated via allosteric changes triggered by copper binding. We propose that Cu(II) gains access to this site through a conformational change, promoting disulphide bond cleavage. Electron transfer is likely facilitated by thiol-disulphide exchange^55^, leading to Cu(I) formation. TTM binding impedes the process by forming a stable complex, inducing an allosteric structural rearrangement of domain III.

### Copper Reduction Depends on Disulphide Accessibility Across Globular Proteins

To evaluate whether copper reduction is unique to HSA or a more general property of disulphide-containing globular proteins, we examined three other globular proteins; β-lactoglobulin (BLG), insulin, and lysozyme using XAS and CD spectroscopy (Figure 4a). These proteins vary in disulphide content, flexibility and copper-binding affinity. BLG contains two disulphide bonds and one free thiol^56^, insulin contains three disulphide bonds and no free thiols^57^, and lysozyme contains four disulphide bonds^58^.

**Figure 4.**
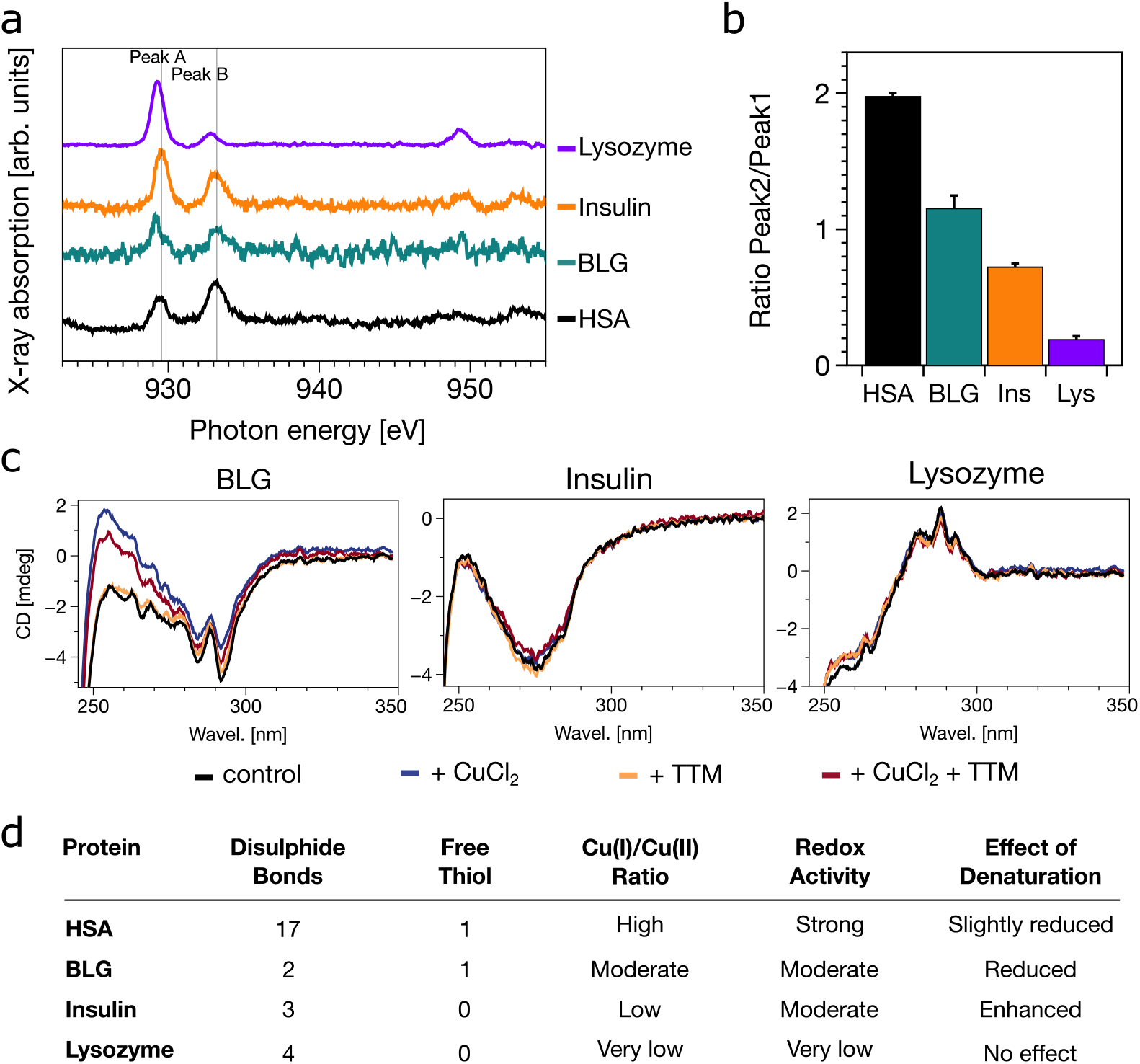
Copper redox activity across globular proteins and correlates with disulphide accessibility and structural flexibility. (a) Soft X-ray absorption spectra at the Cu L_2,3_-edges for HSA (black), β-lactoglobulin (BLG, green), insulin (orange), and lysozyme (purple), each incubated with CuCl_2_ (molar ratio 1:2) at pH 7.4. Cu(I)-associated features are most pronounced in HSA and weakest in lysozyme. (b) Quantified Cu(I)/Cu(II) ratios derived from integrated peak areas of (a). HSA exhibits the highest Cu(I) proportion; lysozyme the lowest. (c) Near-UV CD spectra comparing protein structural changes upon copper and TTM addition. Disulphide sensitive signals (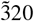 nm) shifts significantly in HSA (Fig 2) and slightly in BLG, and not in insulin and lysozyme. (d) Summary table comparing disulphide architecture, free thiol presence, redox activity, and denaturation sensitivity for each protein. HSA combined high redox activity with flexible disulphide topology; lysozyme, with tightly packed disulphides an no free thiols, shows no redox response.

Soft XAS spectra showed that BLG and insulin weak Cu(I) signals, while the lysozyme showed almost none. CD spectra indicated minor structural rearrangements in BLG and insulin upon Cu or TTM binding, but lysozyme remains unchanged. But none showed an alteration of the disulphide bonds. In general, HSA showed the strongest binding affinity to copper, while lysozyme exhibited the lowest^35,59–62^.

Notably, heat denaturation abolished Cu(II) reduction activity, indicating that native fold is essential. Heat-induced denaturation leads to the loss of native structure and the formation of large amorphous aggregates^63^. As a result, all proteins display reduced Cu-binding capacity. This may be due either to disruption of key disulphide bond positioning or to loss of allosteric communication required for redox activation. Among the proteins tested, only HSA exhibited robust redox modulation with associated conformational rearrangements. Insulin’s reduced Cu(II) affinity upon dimerisation has been previously documented^61^. This highlights the importance of both disulphide topology and structural flexibility in enabling protein-driven copper redox activity.

### Disulphide Cleavage in SOD1 Enhances Copper-Cysteine Interactions and Facilitates Redox Modulation

To validate the role of disulphide bonds in copper redox modulation, we studied superoxide dismutase 1 (SOD1) as a comparative model. SOD1 requires a bound copper ion, which can cycle between two oxidation states, enabling its function^59,64^. Soft XAS measurements across a mildly acidic pH range (pH 5.0-pH 7.4) showed stable Cu(I)/Cu(II) ratios, indicating that histidine protonation alone does not significantly affect copper oxidation state (Fig. 5). This aligns with our observations in HSA, where redox activity was linked instead to disulphide bond dynamics.

**Figure 5.**
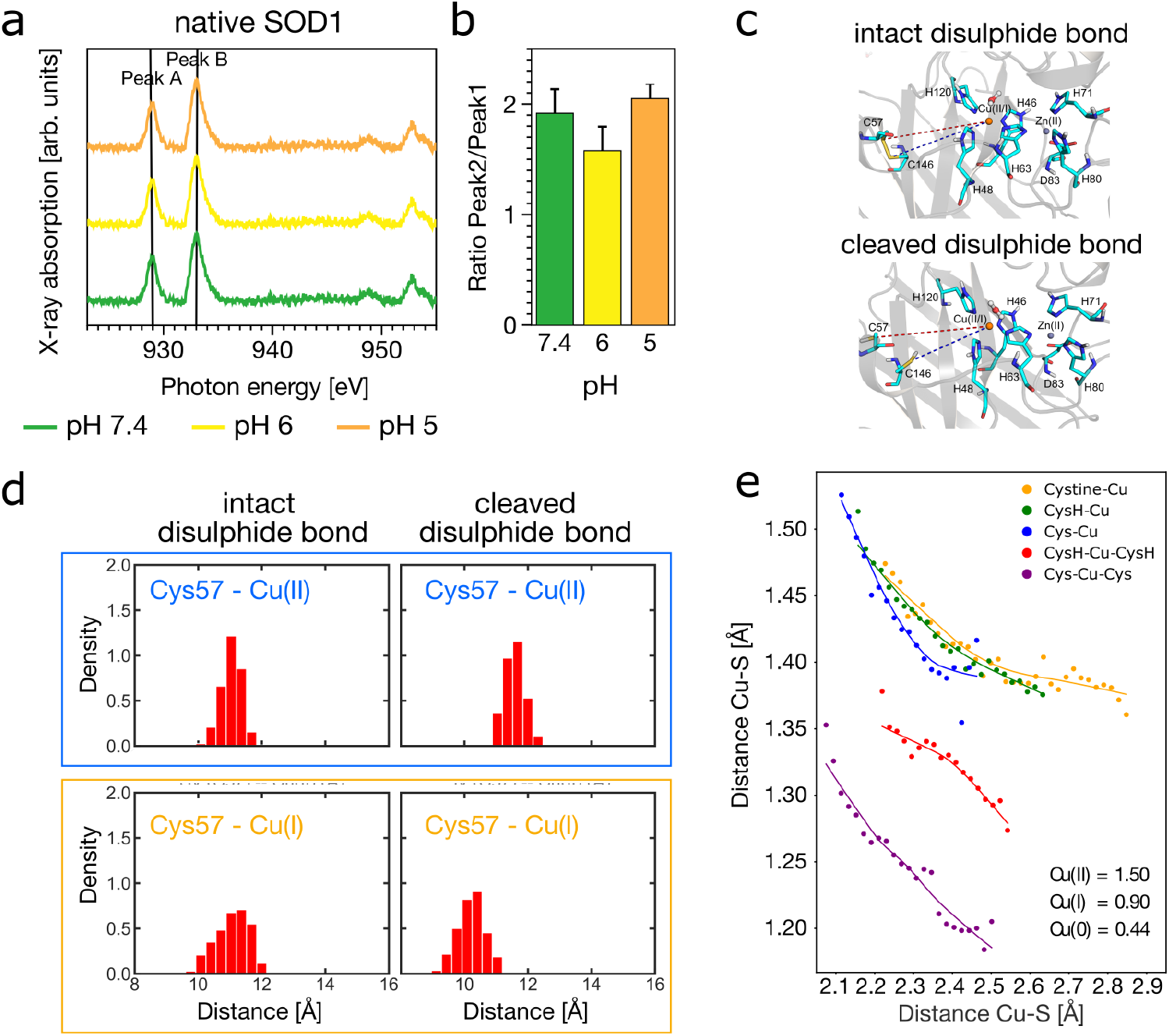
Disulphide bond cleavage in SOD1 enhances Cu-thiol interaction and facilitates Cu(II) reduction. (a) Soft XAS spectra of native SOD1 at pH 7.4 (green), 6.0 (yellow), and 5.0 (orange), showing consistent Cu(I)/Cu(II) ratio, indicating that histidine protonation alone does not modulate copper redox state. (b) Ratio of Cu(I) to Cu(II) peak areas extracted from (a), confirming stable redox state across pH conditions. (c) QM regions used in QM/MM MD simulations, comparing intact (top) and cleaved (bottom) Cus57-Cys146 disulphide bond in SOD1. (d) Histograms showing Cu-Cys57 distances from QM/MM MD simulations. Disulphide cleavage shortens the Cu-S distances, especially in the Cu(I) state, enhancing Cu-thiolate coordination. (e) DFT-calculated Mulliken charges on copper in different ligand environments. Coordination by two deprotonate cysteines (2xCys) significantly reduces copper charge, consistent with Cu(I) formation. In contrast, cystine or neutral cysteine (Cys-H) do not support this reduction.

Classical MD simulations showed that in native (oxidised) SOD1, Cu remains distant from redox-active cysteines (Cys57/Cys146) (Fig. S10). However, upon disulphide bond cleavage, Cu(II) migrates closer to Cys57, suggesting in-creased Cu-S interaction. QM/MM and DFT calculations further revealed that, while Cu(II) remains stable when coordinated by cystine or neutral thiol, coordination by two deprotonated thiolates results in a significant drop in charge density on copper (consistent with Cu(II) reduction) (Fig. 5, S11).

These findings support a model in which disulphide bond cleavage enables Cu-thiolate interaction and electron transfer. The same mechanism likely operates in HSA at Cys392-Cys438, suggesting that disulphide-gated copper redox activity is a generalizable feature in globular proteins.

## Discussion

Transition metals are essential to many biological processes, from energy production to oxidative stress response, and intracellular signalling. Copper, in particular, has critical roles in neurotransmitter synthesis and oxidative stress regulation, but disruptions in copper homeostasis can lead to neurotoxicity and disease. Traditionally, protein-mediated Cu(II) reduction has been associated with pathological or unfolded proteins such as amyloid proteins, mediated primarily by His, Trp, and Cys residues^19,65,66^. However, our findings demonstrate that well-folded globular proteins like HSA also possess intrinsic Cu(II)-reducing capability, independent of classical reducing agents.

HSA, the most abundant protein in plasma, is a primary copper transporter. While only a small fraction of circulating HSA typically carries copper^67^, this increases in disorders such as Wilson disease^68^, where copper-bound HSA contributed to endothelial damage and blood-brain barrier disruption^69^. Previous studies have shown that HSA can bind both Cu(II) and Cu(I)^48^, the latter usually formed via chemical reduction leading to a digonal conformation^28,48^. Here, we show that Cu(I) also forms in HSA without external reductants, and exhibits distinct coordination from ascorbate-reduced Cu(I), suggesting distinct intrinsic reduction site.

As a first thought, the involvement of the conserved residue Cys34, the only free thiol in HSA, which constitutes the largest pool of reactive thiols in plasma, appears to be the most obvious. Its thiol group, typically reduced^70^, is essential to protect against oxidative stress, which can lead to a significant increase in oxidized Cys34 (from 35% to 70%^71^). Cysteines are efficient Cu(II) reductants^8,9^ and important low molecular mass chelator in blood plasma^10^. Apart from methionine residues, cysteines are the most common amino acid in proteins, with abundances in mammals of approximately 2.03 to 2.45%^72^, affecting the protein structure, stability, and function^73–75^. Although Cys34 is a likely candidate for redox activity, our SAXS and structural data indicate that there are no significant conformational changes in domain I, where Cys34 and the ATCUN motif reside. Furthermore, the ascorbate-reduced Cu(I) is consistent with linear bis-His or His-Cys coordination^48^, but the Cu(I) formed intrinsically by HSA appears to adopt a four-coordinate geometry, potentially including Cys392, Cys438 and His440, Arg445 and Lys389. This points to a distinct site of reduction, structurally and functionally separate from known Cu-binding regions.

Using SAXS-guided modelling and chemical probing, we identified the disulphide bond Cys392-Cys438 in domain III as a likely redox-active site. This bond undergoes substantial lengthening upon Cu and TTM binding, beyond typical disulphide geometry, consistent with bond destabilisation. Reversible thiol modifications, including S-cysteinylation of Cys392 and Cys438, have been observed in HSA from hyperlipidaemia patients^76^ and hyperactivity could be observed^77^, supporting their potential involvement in redox activity. QM/MM and DFT simulations support the hypothesis that disulphide bond cleavage facilitates Cu-thiolate interaction and electron transfer, leading to Cu(II) reduction. These data suggest a redox switching mechanism gated by disulphide accessibility.

TTM, a copper chelator and therapeutic agent, interferes with this mechanism. While it stabilises HSA thermally and induces structural compaction, it also alters redox activity and structural dynamics in domain III. The conformational closure observed between domain I and III might disrupt access to the Cys392-Cys438 region, thereby inhibiting copper reduction. These findings imply that TTM may modulate not only Cu bioavailability but also protein structure and function, potentially affecting ligand binding at known pharmacological sites, i.e. by closing the gap between the subdomain IA-IB-IIA and IIB-IIIA-IIIB.

Comparison with other globular proteins reinforces the specificity of the HSA mechanism. β-lactoglobulin (BLG) and insulin exhibited modest redox activity, while lysozyme does not reduce Cu(II) at all. This variability is correlated with differences in disulphide topology and structural flexibility. Denaturation experiments show that folded structure is essential for Cu reduction, supporting the idea that specific protein architecture enables functional disulphide-mediated redox switching. HSA mediated reduction remained relatively persistent even after thermal denaturation, which may be attributed to the high stability of specific disulphide bonds within HSA^34^

We further validated the mechanism in SOD1, a classical metalloprotein with a known Cu site and internal disulphide bond (Cys57-Cys146). XAS data showed stable Cu redox state over a pH range, ruling out histidine protonation as a driver. QM/MM simulations revealed that disulphide cleavage enhances Cu proximity to Cys57, and DFT calculations confirmed that thiolate coordination lowers Cu charge, consistent with Cu(I) formation. This supports a model where disulphide dynamics regulate copper redox state though site accessibility and coordination changes.

Taken together, we propose a general mechanism in which high-affinity Cu(II) binding initiates structural changes that expose a disulphide-containing secondary site. Cleavage of this bond permits Cu-thiolate coordination and facilitates electron transfer. This two-site model, exemplified by both HSA and SOD1 (see Fig. 6), suggests that redox regulation by disulphide bonds may be a broader feature of globular proteins than previously appreciated.

**Figure 6.**
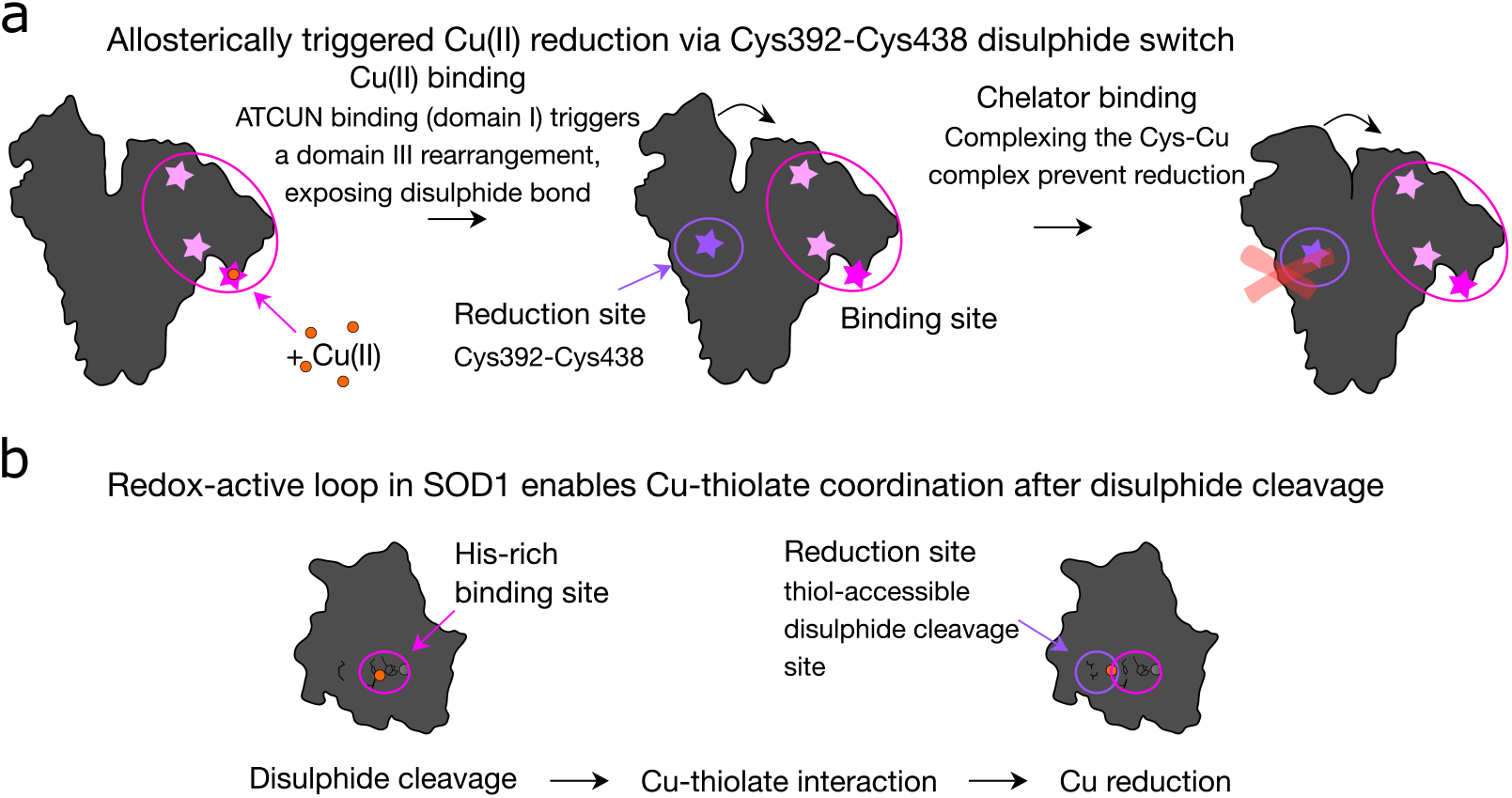
Mechanistic model of copper reduction in globular proteins via disulphide-gates redox switching. (a) In HSA, Cu(II) binds to the high-affinity ATCUN motif (domain I), triggering an allosteric structural rearrangement in domain IIIA/B. This exposes the disulphide bond Cys392-Cys438, enabling thiol coordination and electron transfer. The Cu(II) is subsequently reduces to Cu(I) in a four-coordinate geometry. Chelation by tetrathiomolybdate (TTM) prevents this reduction by interfering with conformational flexibility and disulphide accessibility. (b) In SOD1, Cu(II) is initially bound at the histidine-rich active site. Cleavage of the intramolecular disulphide bond (Cys57-Cys146) enables Cu to shift towards Cys57, enhancing thiolate interaction and facilitating Cu(I) formation. This supports a dynamic redox loop that is modulated by structural integrity and disulphide status.

These findings expand our understanding of protein-based redox chemistry and copper biology. By uncovering a structural mechanism for Cu(II) reduction in folded proteins, this work opens new directions for studying copper trafficking, allosteric control, and redox regulation in physiological and pathological settings. Further research may explore whether other disulphide-containing plasma proteins employ similar redox switches, and how these mechanisms interface with disease states involving copper imbalance.

## Conclusion

Our results uncover a novel copper redox mechanism in globular proteins driven by structural rearrangements and disulphide bond dynamics. In HSA, high-affinity Cu(II) binding initiates allosteric changes that expose a reactive disulphide site, enabling electron transfer and Cu(I) formation. This mechanism is impaired by chelators like TTM, and absent in proteins with inaccessible disulphides or rigid structures. Validation in SOD1 supports the generality of disulphide-gated copper reduction. These findings broaden the known scope of redox activity in plasma proteins and suggest new avenues for exploring copper regulation in health and disease.

## Methods

### Sample preparation

Proteins used in this study, e.g. HSA, BLG, insulin and lysozyme were purchased from Sigma-Aldrich A3782, L2506, I2643 and L1667, respectively. The proteins were dissolved in 20 mM HEPES buffer, pH 7.4 at a concentration of 1.5 mM and subsequently purified using a Superdex 75 increase on a NGM system (Biorad). The same buffer was used as a running buffer. The high concentrated samples were aliquoted, frozen with liquid nitrogen, and stored at -20^*°*^C until further use. The SOD1 protein was expressed and purified by established protocols^78^. Briefly, SOD1 was expressed in BL21(DE3)pLysS competent cells and after harvesting the cells were lysed by sonication (3x 1 min on ice, 1 s on, 0.5 s off). The cleared lysate was incubated at 65^*°*^C for 30 min in a water bath and centrifuged. The SOD1 was precipitated with ammonium sulphate in three steps (50%, 60% and 90%) and subsequently purified using a HiLoad Superdex75 16/60 column and Hitrap Q FF column. DAH-NH2, DTHFPI-NH2, and MEHFPGP-NH2 were synthesized at the Institute of Technology of the University of Tartu, Estonia. The peptides were synthesized on an automated peptide synthesizer (Biotage Initiator+ Alstra, Sweden) using a fluorenylmethyloxycarbonyl (Fmoc) solid-phase peptide synthesis strategy and purified by reverse-phase liquid chromatography on a C4 column (Phenomenex Jupiter C4, 5 µm, 300 A, 250 × 10 mm) using a gradient of acetonitrile/water containing 0.1% TFA. The molecular weight of the peptides was determined by a MALDI-TOF mass spectrometer (Bruker Microflex LT/SH).

### Soft X-ray absorption spectroscopy

The samples for soft XAS measurements were diluted to a final concentration of 100 µM, supplemented with 200 µM CuCl_2_ and in the samples to investigate the chelating mechanism of TTM additional 100 µM TTM for a final molecular ratio of 1:2:1 (protein:CuCl_2_:TTM). The samples were incubated for 1 to 2 h at room temperature and deposited on the surface of a silicon dioxide wafer. The deposited samples were immediately dried with nitrogen gas to form a thin film. Protein samples denoted as heat denatured, were incubated 80^*°*^C for 30 min before CuCl_2_-addition. Additional metal ions were not added to SOD1 prior measurement. SOD1 was purified according to established protocols^78^. After purification, SOD1 was diluted to 100 µM and immediately deposited on a silicon dioxide wafer or incubated with lipids DOPC:DOPG (4:1) for 1 h at 37^*°*^C (lipid/SOD1 ratio: 15/1) before drying. The lipid vesicles were prepared by dissolving 11.25 mg/ml 1,2-dioleoyl-sn-glycero-3-phosphocholine (DOPC), 1,2-dioleoyl-sn-glycero-3-phospho-(1’-rac-glycerol) (DOPG) (4:1) in 100 mM NaCl solution. The lipids were dried with nitrogen gas on a glass bottle surface and then with a desiccator for 2-3 hours. Afterwards 100 mM NaCl solution was added, vortexed for 5 min and then sonicated for 15 min to generate lipid vesicles with smaller and homogeneous size.

The XAS spectra were recorded at the X-Treme beamline^79^ at the Swiss Light Source, Paul Scherrer Institut, Switzerland. To protect the samples from beam damage, an attenuated photon flux and a defocused X-ray spot (0.5 mm^2^) on the sample were chosen. Specifically, the impinging X-ray photon flux per area was 0.06 photons/sec/nm^2^ at the Cu L_2,3_-edges. The spectra were recorded over a timescale of several tens of minutes and when changes were observed, the measurement time was reduced and fresh sample spots were recorded for repeats. The spectra were recorded at room temperature in normal incidence of the X-ray beam in the total electron yield mode (TEY) using on-the-fly scanning. Spectra were normalized for the sum of the peak areas to yield a constant of 1.0 after subtracting the baseline. Reference spectra at the Cu L_2,3_-edges were obtained on a drop-cast film of Cu(II)-phthalocyanine on silicon dioxide wafer.

### Hard X-ray absorption spectroscopy

Hard X-ray absorption spectroscopy in fluorescence mode was conducted at the SuperXAS beamline at the Swiss Light Source, Paul Scherrer Institut, Switzerland. The samples were prepared and analysed in aqueous solution with a protein ratio of 1:2 to CuCl_2_. The final CuCl_2_ concentration was 1 to 1.5 mM. After an incubation of 1 to 2 h, the samples were loaded into the sample holder and flash frozen with liquid nitrogen. The XANES of the Cu K-edge was recorded using a cryoholder. 5 spectra per sample were recorded and analysed using the XAS data processing tool Athena.

### MD simulation

#### Two Cys residues with copper ions in the SOD1 structural model

In order to explore the dynamics between the two CYS residues with copper ion, four systems were set up for the SOD1 as structural model. One is for SOD1/Cu(II) with and without a disulphide bond between C57 and C146 (referred as SOD1/Cu(II)/SS and SOD1/Cu(II)/SHSH), the other one is SOD1/Cu(I) with and without a disulphide bond between C57 and C146 (referred as SOD1/Cu(I)/SS and SOD1/Cu(I)SHSH). The crystal structure of SOD1 (PDB ID: 1HL5^80^, chains A and H) was used as the starting structure for all simulations of the four systems reported in this work. The simulations were performed at pH 7.0. The protonation states of His were chosen based on hydrogen bond network and ions coordination manually examined, H46, H71 and H80 are protonated at N*ε*, H48 and H120 are protonated at N*δ*, while H63 is double deprotonated as it is coordinated with both Zn^2+^ and Cu^2+^. All Asp and Glu residues were negatively charged while all Arg and Lys residues were positively charged. The protein was immersed in a water box with a 10 Å buffer of TIP3P water molecules^81^, and Na^+^ and Cl^−^ were added to neutralize the system and maintaining 0.15 mol/L NaCl mimicking the physiological condition. The protein was described using the Amber ff14SB force field, while the Lennard-Jones 12-6-4 potential of Zn^2+^ and Cu^2+82^ against TIP3P water model were used [ref: Li]. The LEaP module of AmberTools 24^83^ was used to generate the topology and coordinate files for the classical MD simulations, which were carried out using the CUDA version of the PMEMD module of the AMBER 24 simulation package^84^. The solvated system was first subjected to 5000 steps steepest descent minimization, followed by 5000 steps conjugate gradient minimization with positional restraints on all heavy atoms of the solute, using a 50 kcal ^−1^ Å^−2^ harmonic potential. The minimized system was then heated up to 300 K using the Berendsen thermostat, with a time constant of 1 ps for the coupling, and 50 kcal ^−1^ Å^−2^ positional restraints applied over three 500 ps steps of heating process. The positional restraints were then gradually decreased to 5 kcal mol^−1^ Å^−2^ over five 500 ps steps of NPT equilibration, using the Berendsen thermostat and barostat to keep the system at 300 K and 1 atm. For the production run, each system was subjected to 100 ns of sampling in an NPT ensemble at constant temperature (300 K) and constant pressure (1 atm), controlled by the Langevin thermostat, with a collision frequency of 2.0 ps^−1^, and the Monte Carlo barostat with a coupling constant of 1.0 ps. The SHAKE algorithm^85^ was applied to constrain all bonds involving hydrogen atoms. A cut-off of Å was applied to all non-bonded interactions, with the long-range electrostatic interactions being treated with the particle mesh Ewald (PME) approach^86^. A time step of 2 fs was used for all the classical simulations, and coordinates were saved from the simulation every 5 ps. Three independent runs were performed.

#### Copper interaction with model proteins

Molecular dynamics simulations were performed to investigate Cu(I)/Cu(II) interactions with model proteins. The simulations were initiated using the crystal structures of Albumin (PDB ID: 7wlf), β-lactoglobulin (PDB ID: 3npo), and Lysozyme (PDB ID: 1rex). The systems were prepared and solvated using the CHARMM-GUI solution builder module^87^. Potassium ions (K^+^) were added to achieve a final concentration of 150 mM, while copper ions (Cu(I) or Cu(II)) at 50 mM were introduced to facilitate protein interaction. Chloride ions (Cl^−^) were included to neutralize the systems. TIP3P water molecules were added, ensuring a cubic simulation box with an approximate edge length of 85 Å. Molecular dynamics (MD) simulations were performed with CHARMM FF using GROMACS 5.0^88^. The MD integrator was used with a time step (Δ t) of 2 fs. The total simulation time consisted of 100,000,000 steps, corresponding to 0.2 µs of simulation time. Trajectory coordinates, velocities, and forces were saved every 1 ns (500,000 steps). Energy and log files were recorded at 2 ps intervals (1,000 steps). A Verlet cut-off scheme was applied with a neighbour list update frequency of 20 steps. For van der Waals interactions, we applied a simple cut-off method with a force-switch modifier from 1.0 nm and 1.2 nm. Long-range Electrostatic interactions were computed using the Particle Mesh Ewald (PME) method with a cut-off of 1.2 nm for the real-space component. Temperature coupling was achieved using the Nose-Hoover thermostat with separate temperature groups for solute (SOLU) and solvent (SOLV). A temperature of 303.15 K was maintained with a relaxation time (*τ*) of 1 ps. Isotropic pressure coupling was implemented using the Parrinello-Rahman barostat, targeting a pressure of 1.0 bar with a compressibility of 4.5×10^−5^ bar^−1^ and a relaxation time (*τ* _*p*_) of 5 ps. All bonds involving hydrogen atoms were constrained using the LINCS algorithm, enabling the use of a 2 fs time step. The centre of mass motion was removed every 100 steps using a linear approach, applied separately to the solute and solvent groups. The system was initialized from equilibrated structures, and the production simulation was continued from the last state of the equilibration phase. Simulation stability was monitored through energy, temperature, and pressure profiles.

#### QM/MM MD simulation

To further access the interplay between the Cu (II/I) and the two CYS residues, quantum mechanics/molecular mechanics (QM/MM)^89^ MD simulations were performed for both of the four systems. ACPYPE^90^ was used to convert the Amber topology and coordinates into the Gromacs format forms. The last snapshot from the classical MD trajectory at simulation time of 100 ns was then used for the subsequent QM/MM MD simulations, which combines Born-Oppenheimer MD simulation, based on density functional theory (DFT), with force-field MD methodology. For SOD1/SS, the QM region consists of the sidechains of H46, H48, H63, H71, H80, D83, H120 and copper as well as zinc ions, resulting in a QM region of 80 atoms (including 7 capping hydrogens), as shown in Fig. S12a. Apart from QM atoms in SOD1/SS, the sidechains of C57 and C146 are added to the QM region for SOD1/SHSH, leading to a total of 92 atoms in the QM region including 9 capping hydrogens (Fig. S12). The QM atoms were centred in a QM box of 18×18×18 Å^3^ for SOD1/SS and 24×24×16 Å^3^ for SOD1/SHSH. All QM/MM MD simulations were performed using Gromacs 2022.5^88^ interfaced with CP2K 2024.1^91,92^, combining the QM program QUICKSTEP^93^ of CP2K and the MD engine of Gromacs. In this code, a real space multigrid technique is used to compute the electrostatic coupling between the QM and MM region. The QM region was treated at the DFT (BLYP) level, employing the dual basis set of Gaussian and plane-waves (GPW) formalism, whereas the remaining part of the system was modelled at the classical level using the same parameters as in the classical MD simulations. The Gaussian triple-*ζ* valence polarized (TZV2P) basis set was used to expand the wave function, while the auxiliary plane-wave basis set with a density cut-off of 400 Ry and GTH pseudopotentials^94,95^ was utilized to converge the electron density. All QM/MM MD simulations were performed under the NVT ensemble at a constant temperature of 300 K using velocity rescaling thermostat with a coupling constant of 50 fs^−1^ and an integration time step of 0.5 fs. First, the system was minimized using the steepest descent method, then it is further equilibrated without any constraint for 5.0 ps. Afterwards, the production runs were performed 56 ps for SOD1/SS and 25 ps for SOD1/SHSH.

#### Ab initio molecular dynamics simulations

Ab initio molecular dynamics (AIMD) simulations for the molecules Cystine, CysH, and Cys in aqueous CuCl_2_ solution were performed with the density functional (DFT) module Q<sc>UICKSTEP</sc>^93^ of the CP2K program package^91,92^ using norm-conserving, scalar-relativistic pseudo-potentials of Goedecker, Teter and Hutter (GTH)^94,95^ and double-*ζ* MOLOPT basis sets^96^. An auxiliary basis set of plane waves was employed to expand the electronic density using an electronic density cut-off of 600 Ry. The exchange and correlation functional of Perdew, Burke and Ernzerhof (PBE) in its original parametrization^97^ augmented by the dispersion correction D3 of Grimme et al.^98^ was employed in all AIMD calculations. The molecules and a CuCl_2_ unit were solvated with 256 water molecules using Packmol^99^ within a sufficiently large simulation cell. Thereafter, a relaxation of the atomic positions and cell parameters was performed (CELL_OPT run). The final configuration was further equilibrated during 3 ps of AIMD within the isobaric-isothermal (NpT) ensemble at 1 bar and 320 K using MD time step of 0.5 fs. The equilibration run was followed by a sampling run of at least 4 ps from which the atomic charges were retrieved by Voronoi integration^100^. The reference Cu charges for Cu(II), Cu(I), and Cu(0) were obtained from simulations of a single Cu atom with two, one, or no Cl atoms in a cell with 256 water molecules, respectively.

### Radius measurement

The radius was determined using the Panta Prometheus (Nanotemper). The samples were prepared to a final concentration 100 µM protein supplemented with different concentration of CuCl_2_, TTM or both using 20 mM HEPES, pH 7.4. The samples were loaded into Prometheus Series capillaries (PR-C002) and the DLS signal was measured.

### Thermodynamic stability

The thermodynamic stability was determined using the scattering as a readout from the Panta Prometheus (Nanotemper). The samples prepared in the same way as for the radii measurements were loaded into Prometheus Series capillaries (PR-C002), closed and heated up from 25^*°*^C to 90^*°*^C with a scanning rate of 1^*°*^C/min.

### CD spectroscopy

CD spectra were recorded at 20^*°*^C using a Chirascan plus with a temperature control Quantum Northwest. Final protein concentration was 100 µM or 200 µM. Differences in secondary structure were first examined, for HSA as an example using PB instead of HEPES, since no influence of CuCl_2_ or TTM on the secondary structure could be observed, the changes in the near-UV were investigated, recording spectra from 250 nm to 350 nm.

### Small angle X-ray scattering

SAXS spectra were collected on the EMBL beamline P12^101^ at Petra III at DESY (Deutsches Elektronen-Synchrotron), Hamburg, Germany. The samples were shipped in liquid nitrogen. Buffer subtraction and data processing were performed using the ATSAS software package, including PRIMUS, to determine the radius of gyration (R_*g*_). Different pdb structures were fitted to the SAXS curves using CRYSOL and flexible refinement using SREFLEX was performed mainly using the models 3b9l (pdb), 4LA0 (pdb), SASDAA6 (SASBDB) and an alphafold model. Since the models lead to rather similar *χ*^2^ values, the presented SREFLEX refinement was performed using model 4LA0 and an alphafold model AF-P02768-F1-v4. 4LA0^51^ is a X-ray structure of HSA complexed with bicalutamide and is one of the few structures containing the AH residues of the ATCUN motif. For the model refinement, the first 30 points were skipped. A consensus structural model was calculated from the best-fitting models based on their x^2^ values. The models were aligned using SUPCOMB and an average model was calculated using PyMOL. The average models were fitted against to the SAXS curves using CRYSOL.

## Acknowledgements (not compulsory)

The synchrotron soft X-ray XAS data was collected at the beamline Xtreme at the SLS, Villigen, Switzerland. The synchrotron SAXS data was collected at beamline P12 operated by EMBL Hamburg at the PETRA III storage ring (DESY, Hamburg, Germany). We would like to thank Melissa Gräwert for the assistance in using the beamline. The synchrotron hard X-ray XANES data was collected at the beamline SuperXAS at the SLS, Villigen, Switzerland. We would like to thank Maarten Nachtegaal and Grigory Smolentsev for assistance in using the beamline. This work was supported by the PSI cross research grant to R.S.H. and the Estonian Research Council grant (PRG 1289) to P.P.

## Author contributions statement

R.S.H., J.L. conceived the experiments, R.S.H., C.L. X.W., H.P., X.S., P.P. conducted the experiments, M.K., Q.L, J.L. performed and analysed computer simulations, R.S.H., C.L. analysed the results, R.S.H., J.L. wrote the manuscript. All authors reviewed the manuscript.

