## Supplementary material for "Protein-Driven Copper Redox Regulation: Uncovering the Role of Disulphide Bonds and Allosteric Modulation"

### Additional information

This Supporting Information provides complementary data and computational analyses that expand on the experimental findings presented in the main manuscript. It includes additional X-ray absorption spectra (Fig. S1, S2) confirming copper oxidation states in HSA and peptide complexes, and extended SAXS and biophysical characterisations (Fig. S3, S4, S5, S6, S7, S8) detailing structural changes, induced by copper and TTM. Fig. S9 extends the XAS analysis to other globular proteins under denaturing conditions, revealing how structural destabilisation influences copper redox behavirour. Figures S10, S11, S12, S13, S14, S15 offer computational insights, including MD simulations, QM/MM and DFT modelling, and AlphaFold-based copper binding predictions, supporting the proposed mechanism of disulphide-gated copper reduction in globular proteins. Together, these data reinforce the structural and redox dynamics underlying copper-protein interactions and provide a broader framework for interpreting redox behaviour across different protein classes.

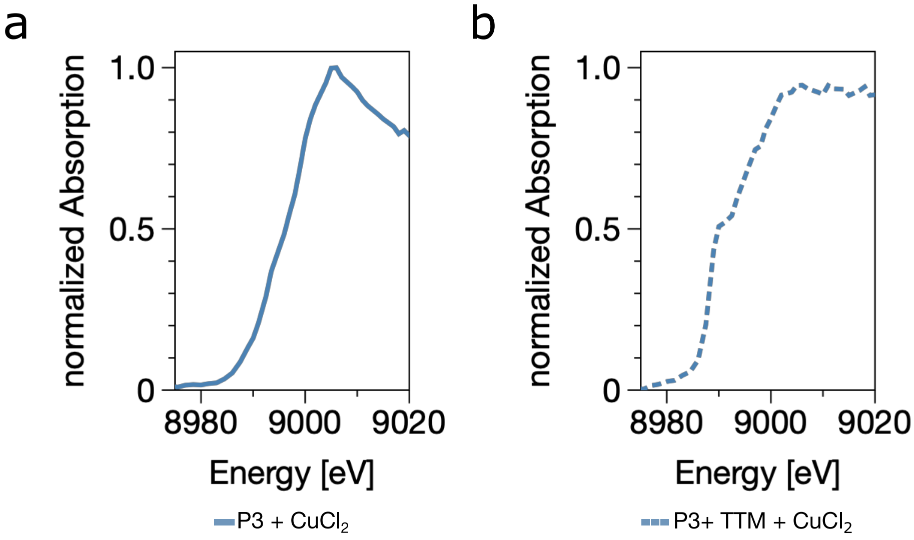


**Figure S1.** Hard x-ray absorption spectroscopy at the Cu K-edge reveals Cu(II) in peptide 3 (MEHFPGP) (a), while the addition of TTM promotes Cu(I) formation (b).

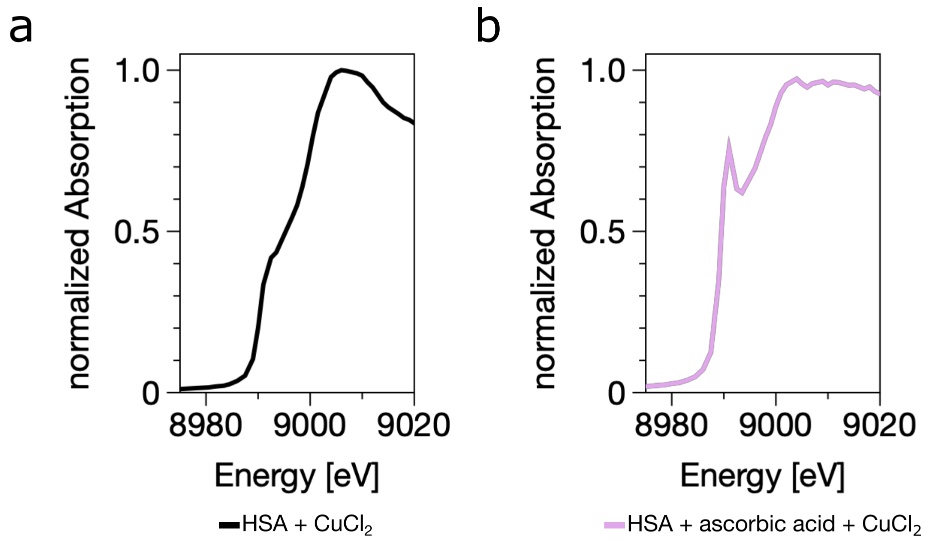


**Figure S2.** Hard x-ray spectroscopy confirms Cu(I) species within HSA (a). However, Cu(I) formed via ascorbate reduction (b) exhibits a distinct coordination geometry, indicating different copper-binding environments.

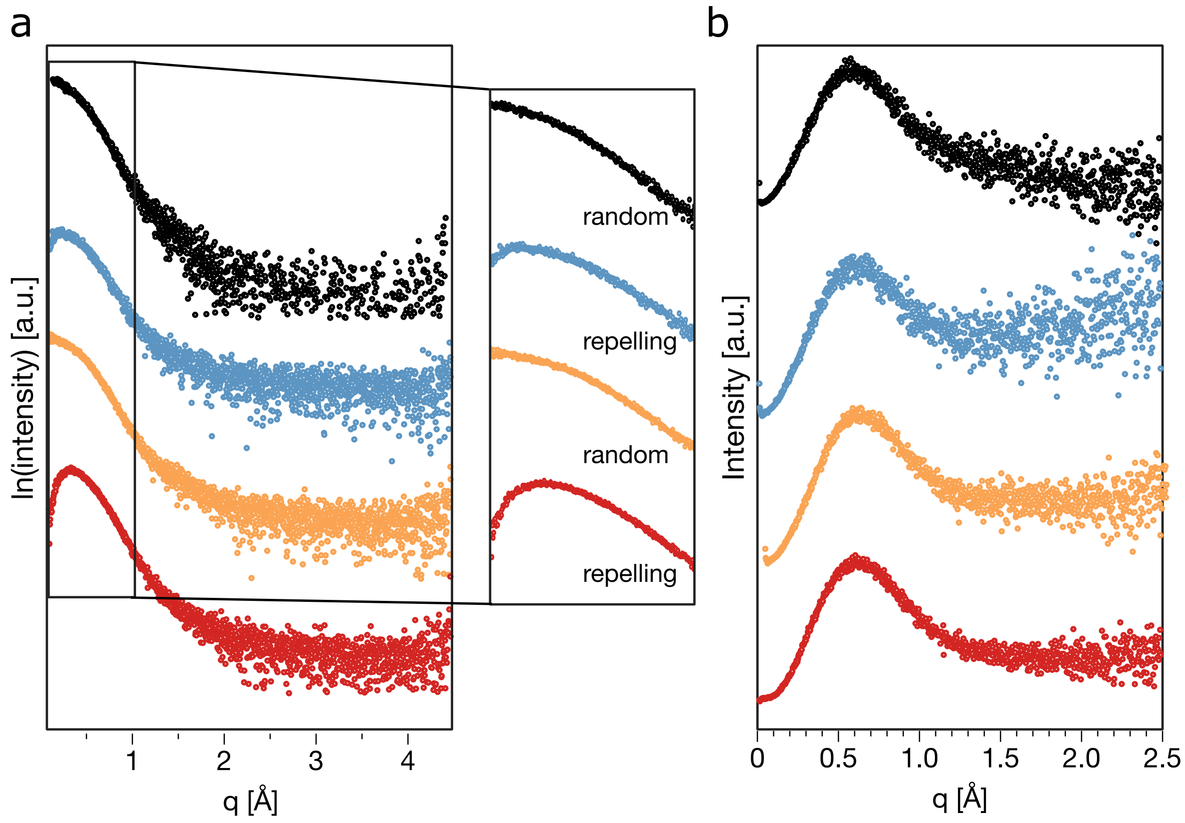


**Figure S3.** (A) Static Small-angle X-ray scattering (SAXS) profiles of HSA (black), with $CuCl_2$ (blue), with TTM (orange) and both (red). Copper induces a repulsive particle distribution; TTM alone does not. (b) Kratky plots indicate that all samples remain folded, though copper increases flexibility slightly.

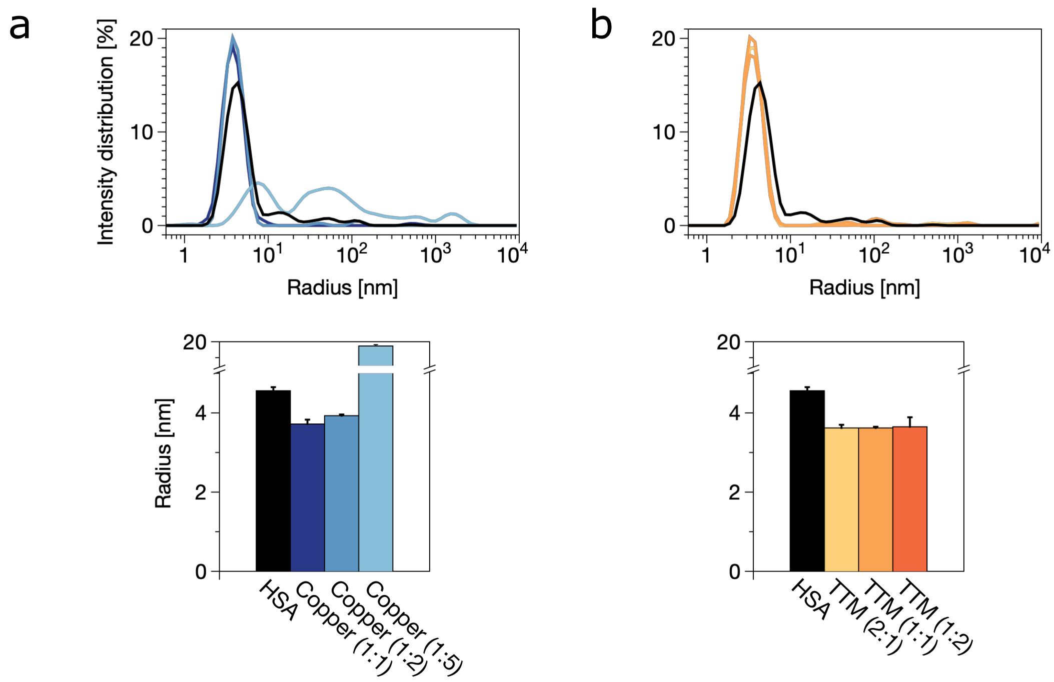


**Figure S4.** Dynamic light scattering (DLS) confirms structural compaction upon addition of (a) $CuCl_2$, and (b) TTM. At higher $CuCl_2$ concentration (1:5), HSA forms larger aggregates.

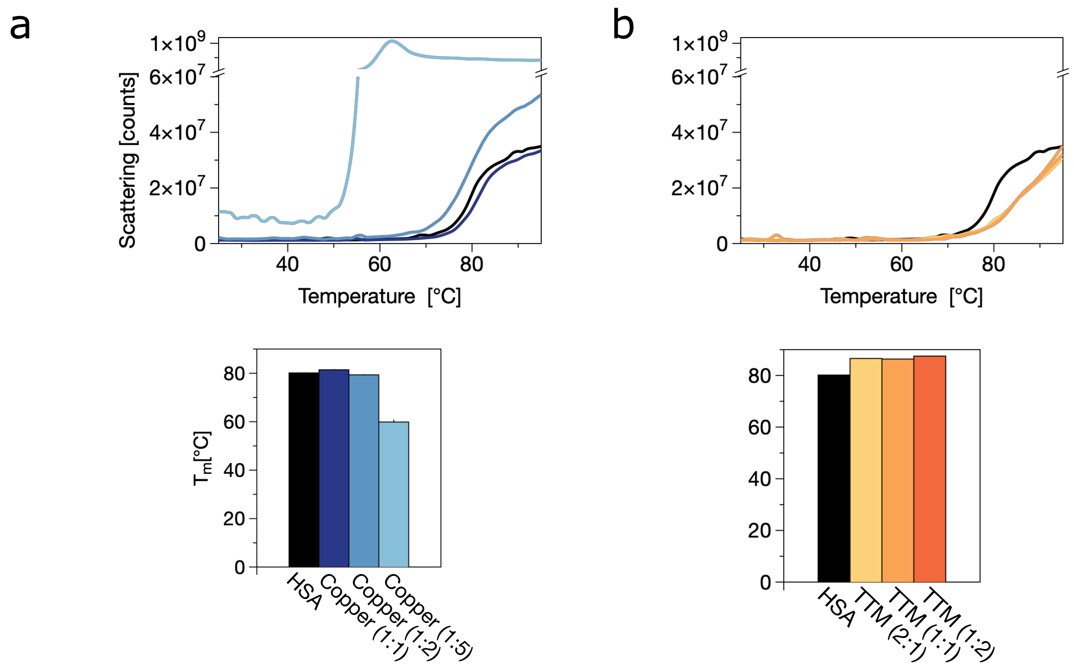


**Figure S5.** (a) Thermal denaturation profiles reveal that high $CuCl_2$ concentrations destabilise HSA and promote aggregation. (b) TTM confers partial thermal protection against denaturation.

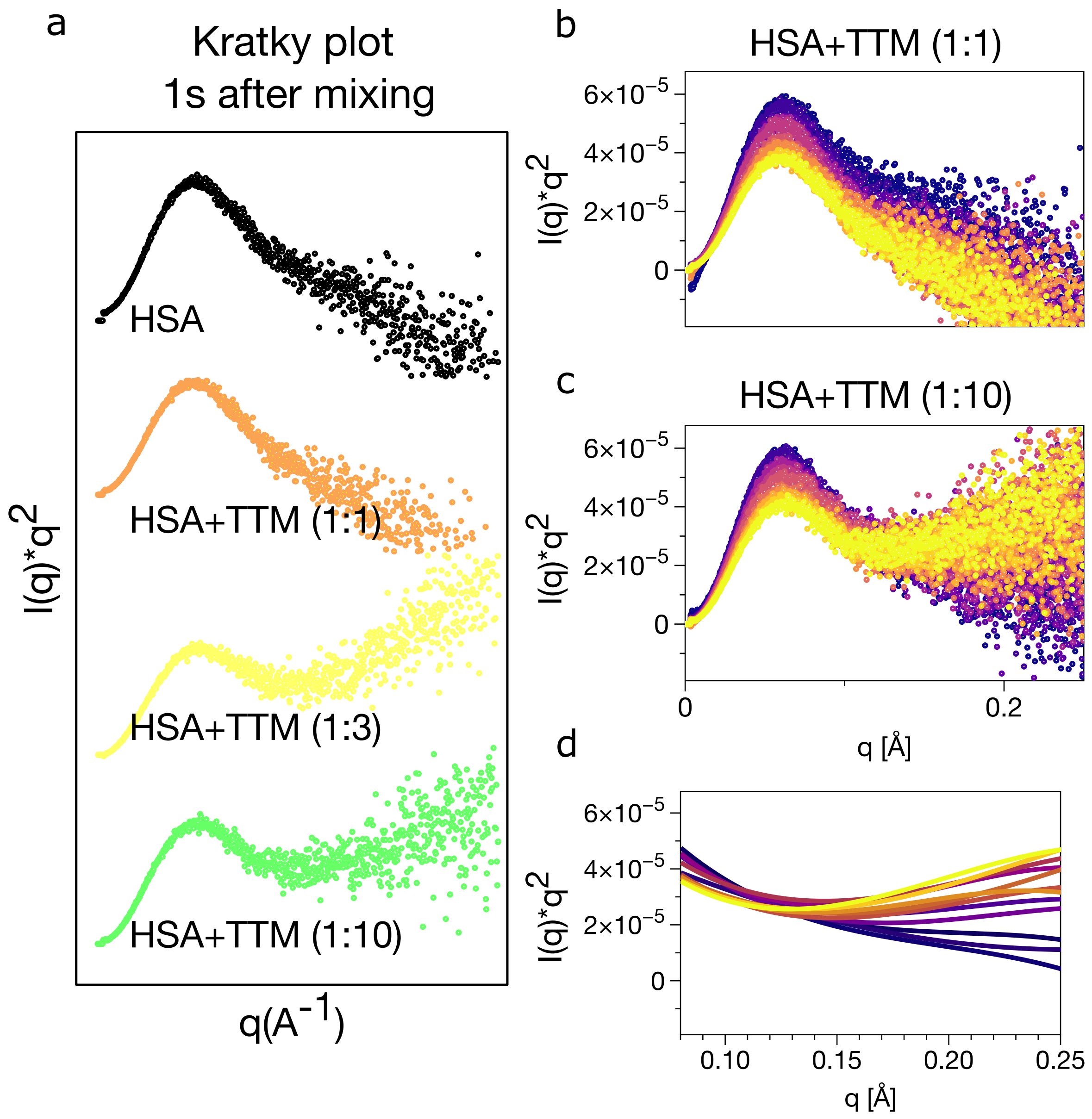


**Figure S6.** Kratky plots from time-resolved SAXS measurement show that (a) HSA and HSA + TTM (1:1) remain folded, while higher TTM concentration (1:3, 1:10) induce partial unfolding. (b, c) Time evolution of the Kratky plots of 1:1 and 1:10 TTM conditions. (d) Cubic fit of HSA+TTM (1:10) shows increased flexibility in the mid-q region (from blue to yellow).

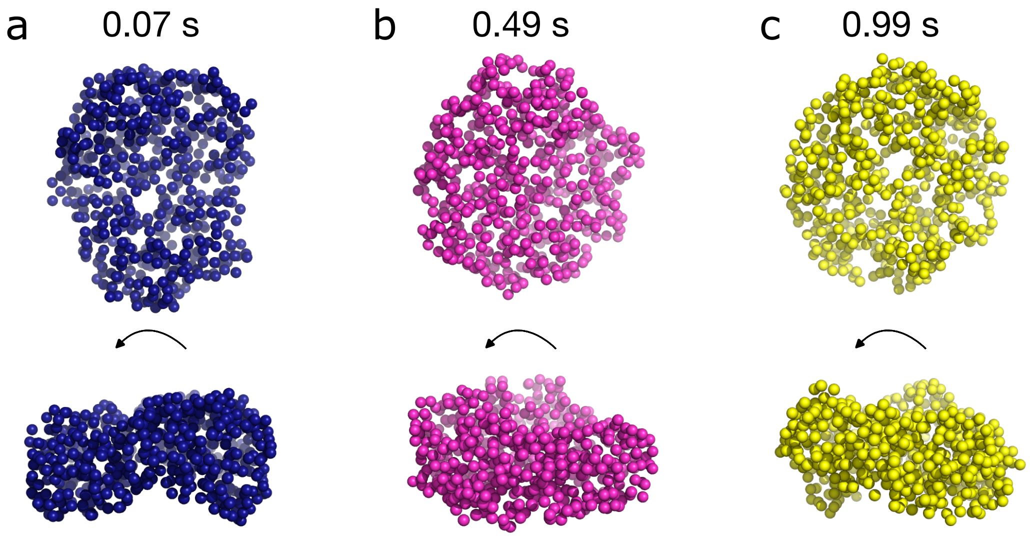


**Figure S7.** Ab-initio GASBOR models of HSA+TTM (1:10 molar ratio) at (a) 0.07 s ($\chi^2$=1.041), (b) 0.49 s ($\chi^2$=1.243) and (c) 0.99 s ($\chi^2$=1.235) after mixing. All models suggest compaction with time under high TTM conditions.

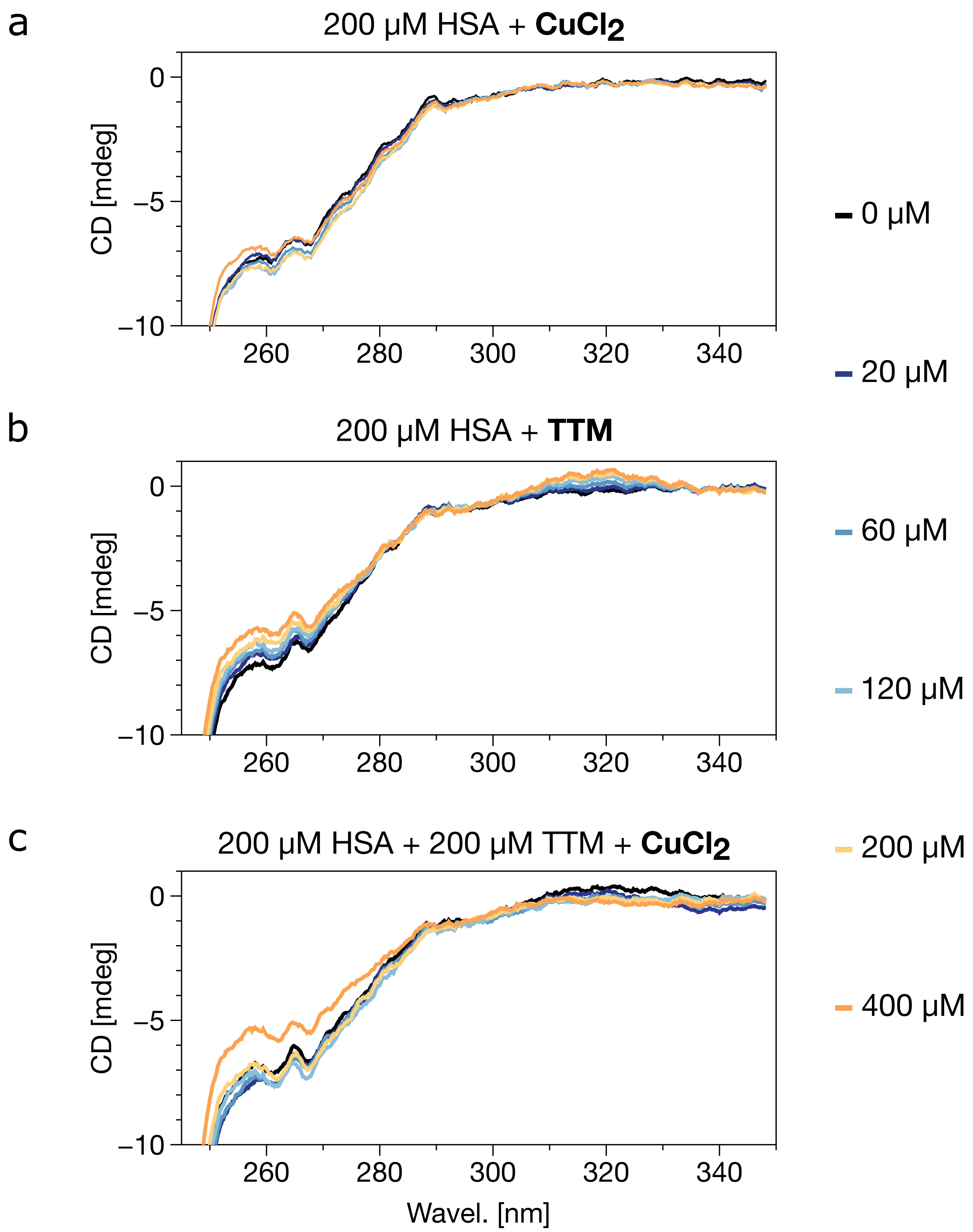


**Figure S8.** Near-UV CD spectra of 200 μM HSA with titration of (a) $CuCl_2$, (b) TTM and (c) $CuCl_2$ to HSA+TTM mixture. The additive was added to a final concentration of 0, 20, 60, 120, 200 and 400 μM. TTM induces changes in aromatic region and a 320 nm S-S-sensitive signal, which disappears upon Cu addition, indicating modulation of disulphide environment.

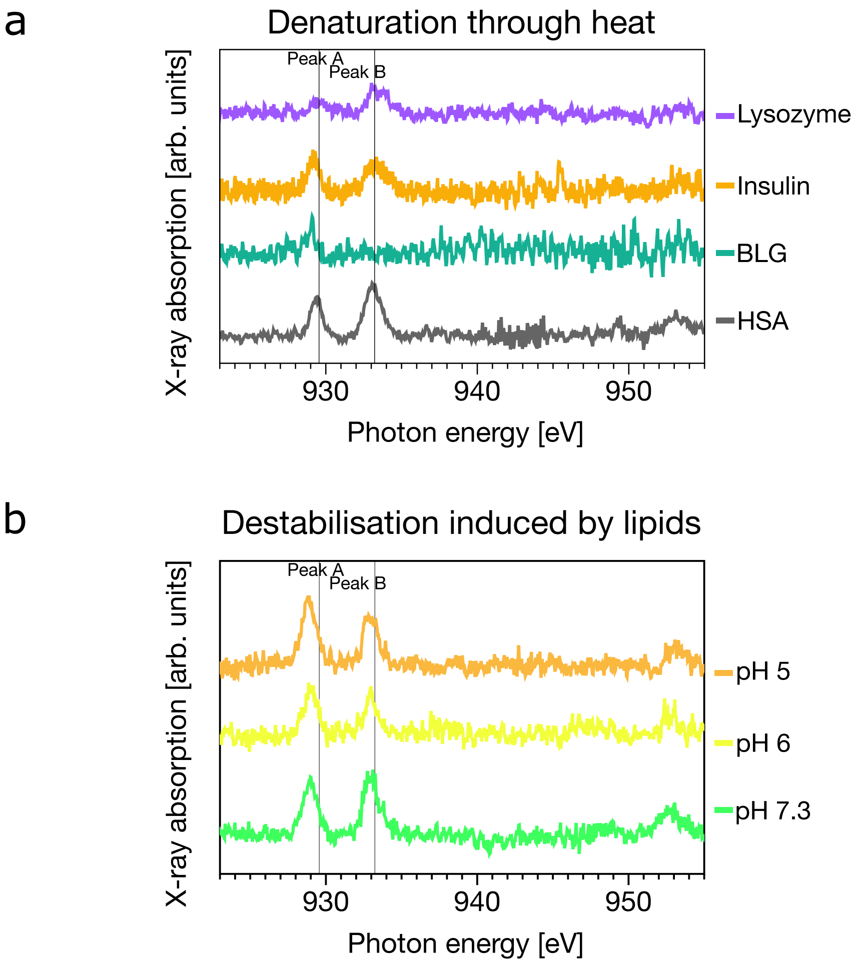


**Figure S9.** (a) Soft XAS spectra of heat-denatured globular proteins. (b) SOD1 spectra after lipid-induces destabilisation. Denaturation affects copper redox behaviour.

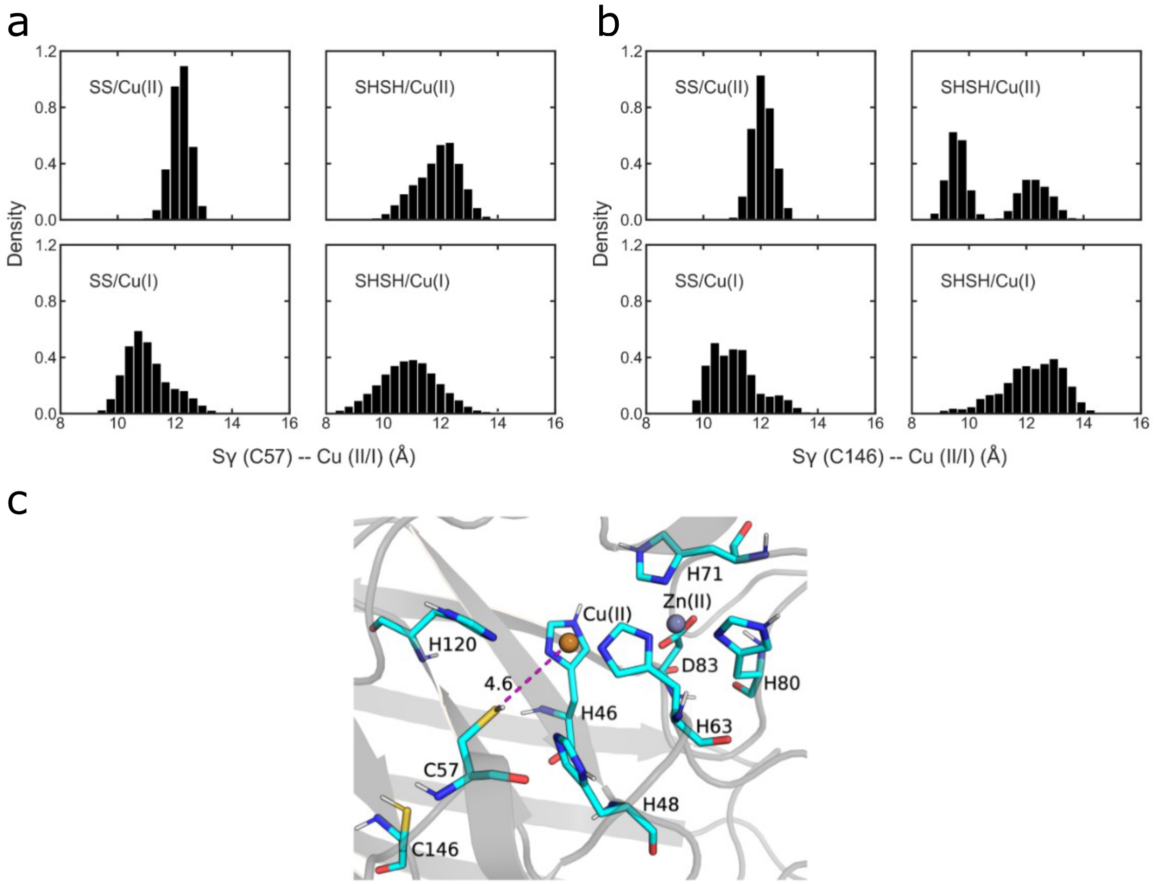


**Figure S10.** Histograms of Cu (II/I) the distances in SOD1 with (a) Cys57 and (b) Cys146 obtained from the classical MD simulations. SS is referred as a disulphide bond between C57 and C146, while SHSH is referred as no disulphide bond between C57 and C146. (c) Disulphide reduction enables Cu to approach 57 more closely.

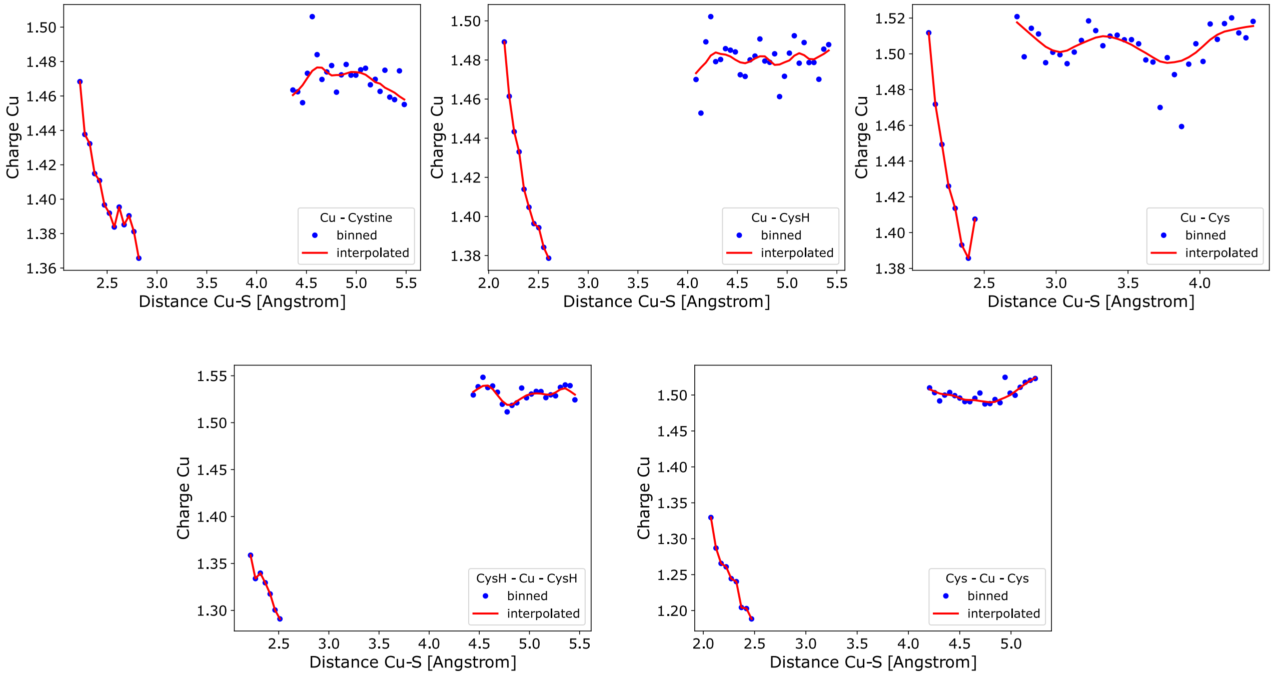


**Figure S11.** DFT-calculated Cu charges in complexes with various cysteine forms. Cu remains in the +2 state with cystine, Cys, or CysH, but is stabilised in the +1 state with two CysH or deprotonated Cys ligands.

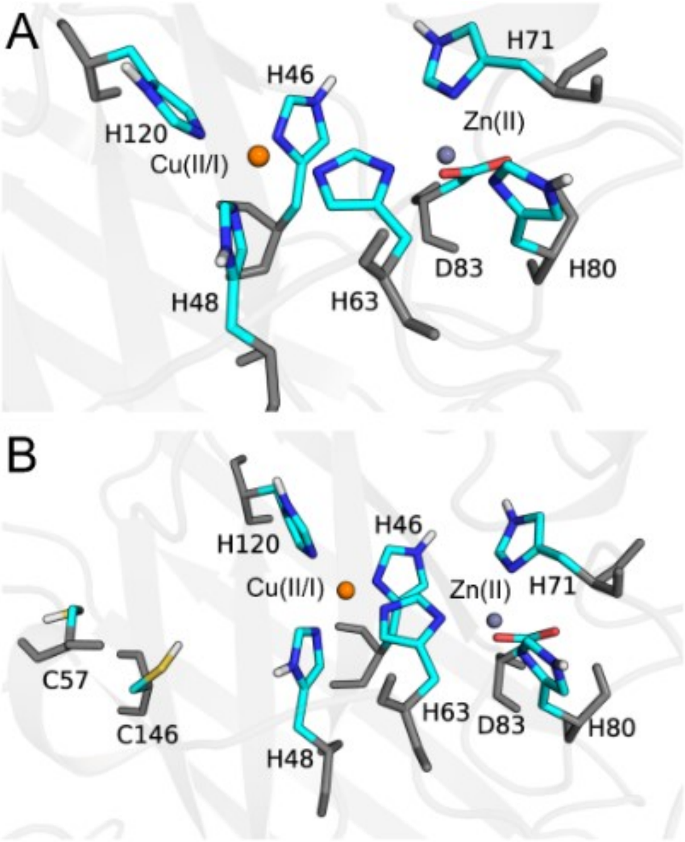


**Figure S12.** QM regions used for QM/MM MD simulations of SOD1 with (a) and without (d) the disulphide bond between Cys57 and Cys146. QM atoms are coloured; others shown in grey (classical force field).

To explore potential copper-binding distributions and conformational flexibility in HSA, AlphaFold predictions and molecular dynamics simulations were conducted. While not experimentally validated in this study, they provide complementary insight into copper–protein interactions and are shown below for reference.

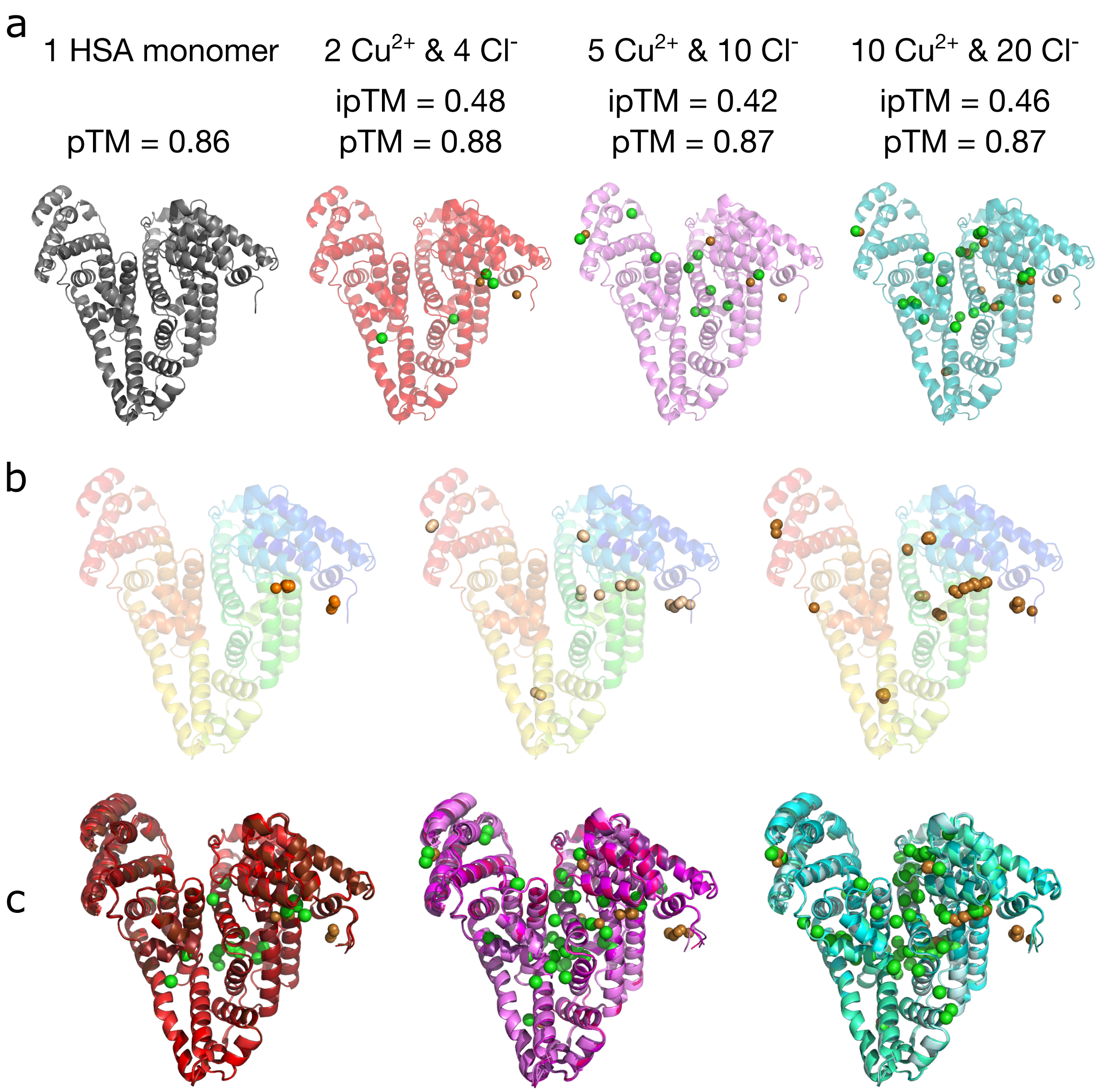


**Figure S13.** AlphaFold-predicted HSA models with varying copper load: (a) model I with the corresponding ipTM and pTM scores; (b) models with 2, 5, and 10 Cu ions (from left to right); (c) overlay of the five predicted models.

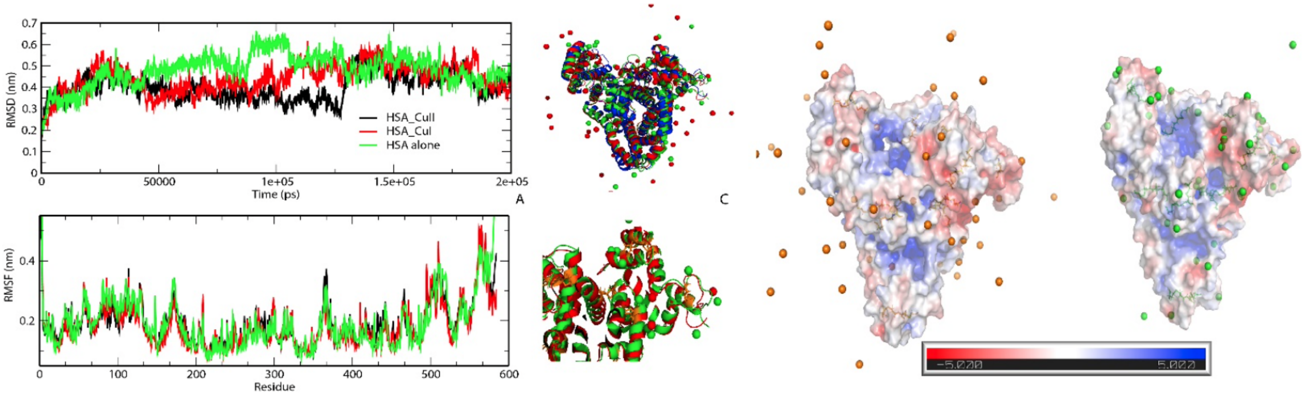


**Figure S14.** MD simulation snapshots showing the Cu(II) and Cu(I) distribution on the HSA surface. Cu(I) remains more surface-associated than Cu(II).

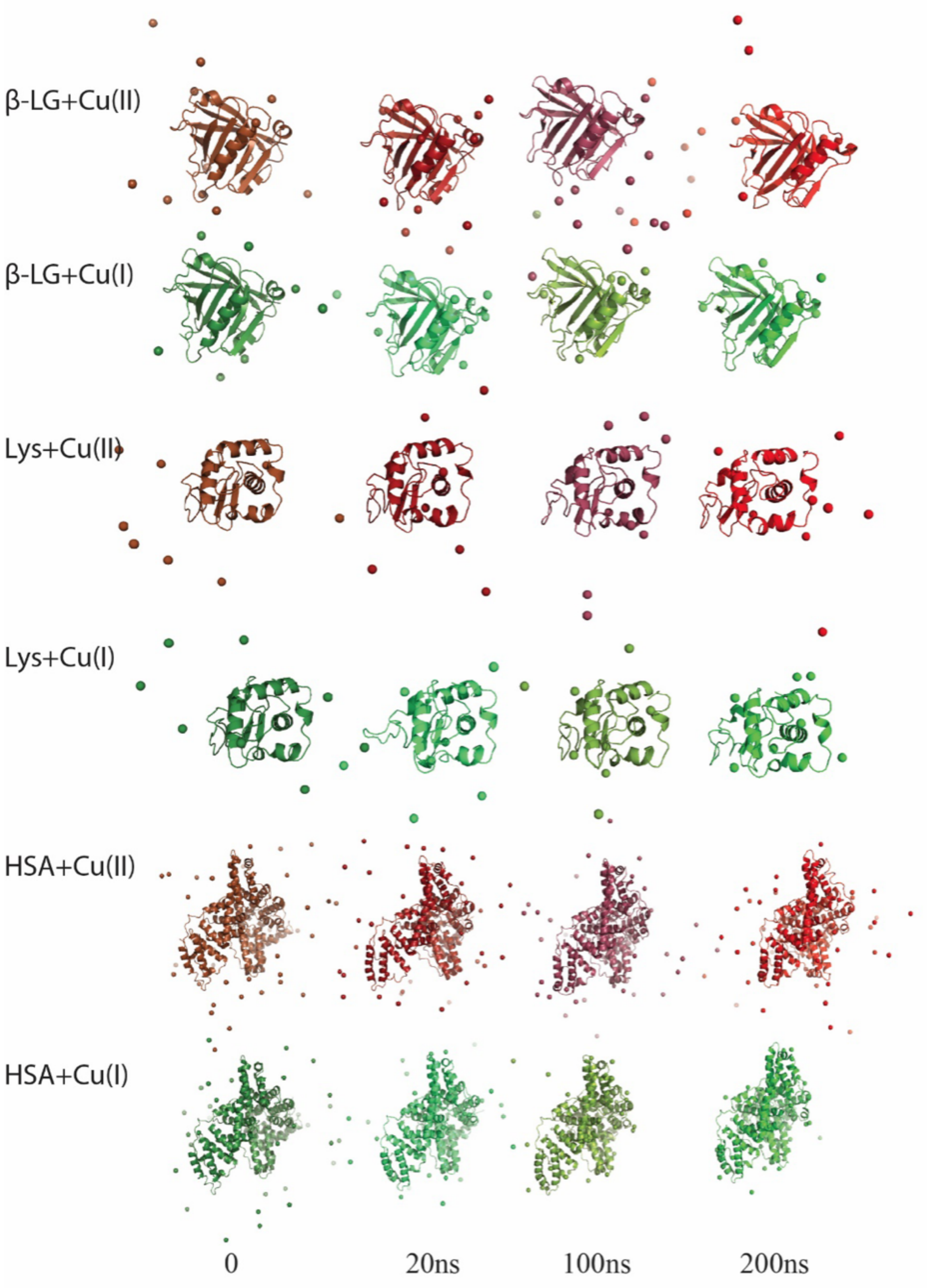


**Figure S15.** Snapshots from MD simulations of multiple proteins in the presence of Cu(II) and Cu(I). Distribution patterns reflex structural features and copper coordination environment.
